# D614G reshapes allosteric networks and opening mechanisms of SARS-CoV-2 spikes

**DOI:** 10.1101/2025.03.07.642081

**Authors:** Fiona L. Kearns, Anthony T. Bogetti, Carla Calvó-Tusell, Mac Kevin E. Braza, Lorenzo Casalino, Amanda J. Gramm, Sean Braet, Mia A. Rosenfeld, Harinda Rajapaksha, Bryan Barker, Ganesh Anand, Surl-Hee Ahn, Lillian T. Chong, Rommie E. Amaro

## Abstract

The SARS-CoV-2 spike glycoprotein binds human epithelial cells and enables infection through a key conformational transition that exposes its receptor binding domain (RBD). Experimental evidence indicates that spike mutations, particularly the early D614G variant, alter the rate of this conformational shift, potentially increasing viral infectivity. To investigate how mutations reshape the conformational landscape, we conducted extensive weighted ensemble simulations of the Ancestral, Delta, and Omicron BA.1 spike strains along the RBD opening pathway. We observe that Ancestral, Delta, and Omicron BA.1 spike RBDs open differently, with Omicron BA.1 following a more direct opening profile until it reaches a “super-open” state wherein it begins to “peel”, suggesting increased S1 flexibility. Via dynamical network analysis, we identified two allosteric communication networks uniting all S1 domains: the established N2R linker and a newly discovered anti-parallel R2N linker. In Delta and Omicron BA.1 variant spikes, RBD opening is facilitated by both linkers, while the Ancestral strain relies predominantly on the N2R linker. In the ancestral spike, the D614-K854 salt bridge impedes allosteric communication through the R2N linker, whereas the loss of this salt bridge in all subsequent VOCs alleviates local frustration and, we believe, accelerates RBD opening. Hydrogen-deuterium mass spectrometry experiments validate these altered dynamics in the D614 region across Ancestral, D614G, and Omicron BA.1 spikes. This study unveils a ‘hidden’ allosteric network, connecting the NTD to the RBD via the 614-proximal region, and the D614G mutation reshapes the fitness landscape of these critical viral glycoproteins.

**Significance Statement:** Our work reveals how the D614G mutation in the SARS-CoV-2 spike protein reshapes its internal communication pathways and speeds up receptor binding domain (RBD) opening, providing mechanistic insight into the evolution and enhanced infectivity of SARS-CoV-2 variants of concern. We also describe differences in opening pathways and relative rates of opening for Delta and Omicron BA.1 spike RBDs relative to the original (Ancestral) coronavirus strain from Wuhan, China.

## Introduction

The SARS-CoV-2 (Severe Acute Respiratory Syndrome COronaVirus 2) viral replication cycle in humans begins with host-cell surface binding and membrane fusion,^1–3^ both steps facilitated by the S, or spike, homotrimeric glycoprotein.^1,4–7^ During expression and viral packaging, the spike protein is folded into a metastable prefusion conformation, with a head region supported in the membrane by a stalk (**Figure 1A)**. Several biophysical events are required to prime the spike for its role as a class I fusion protein.^8^ First, furin must cleave an RRAR recognition site, the furin cleavage site (FCS), splitting the spike sequence into S1 and S2 domains (**Figure 1A,B)**.^9–14^ Second, in its metastable folded state the ACE2 binding motif, known as the receptor binding motif (RBM) within the receptor binding domain (RBD), is highly shielded from recognition by neighboring protein surfaces and by N- and O-linked glycans.^15–19^ As such, this initial “closed” conformation of the prefusion spike must undergo a key conformational change in which at least one of its RBMs emerges from the down/shielded state into an up/exposed state (**Figure 1B)**.^19–21^ Spike conformations with one, two, or three RBDs in the up state are referred to as 1-up, 2-up, and 3-up states respectively. During RBD opening, other domains shift as well including the N-terminal domain (NTD), sub-domain 1 (SD1), sub-domain 2 (SD2), and the fusion peptide proximal region (FPPR) (**Figure 1AB)**.^1,7,9^ Several isolated monoclonal antibodies target key epitopes within spike’s RBD and RBM. As such, all the SARS-CoV-2 vaccines developed to combat the COVID19 pandemic were based on modified spike mRNA sequences or live attenuated viruses with displayed spike glycoproteins.^22^ In fact, it is hypothesized that shielding RBD and RBM epitopes until the spike is near the host cell surface is advantageous to balance infection and replication with immune evasion.^7,8,23^ After RBD opening and ACE2 binding, the S1 domain peels off revealing the S2 core and fusion peptide (FP) (**Figure 1A,B)**.^7,24–26^ Once all three S1 domains are removed, and following another cleavage event to free the FP,^13,14,27^ the prefusion spike structure undergoes massive rearrangement into a postfusion state concomitantly penetrating the neighboring human host-cell membrane in the process.^24–26^ Considering the spike’s key role in host-cell infection as well as its demonstrated value as a vaccine target, much attention has been paid to investigating spike structural biology for the purposes of drug design^28–31^ and pancoronavirus vaccine development.^32–35^

**Figure 1:**
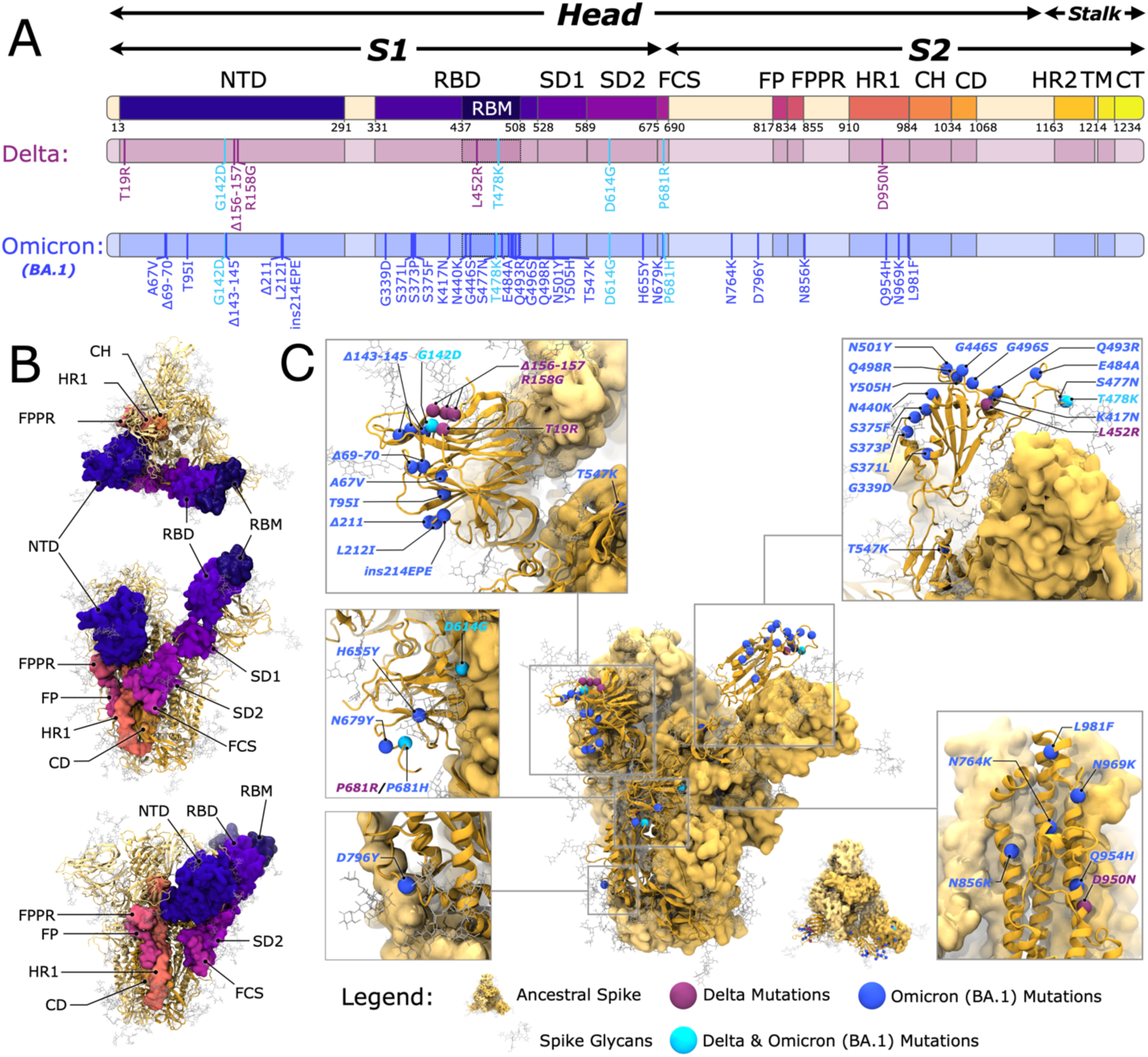
Mutations to the ancestral strain spike protein leading to definitions of Delta and Omicron BA.1 spike proteins considered in this work. **(A)** Definitions of spike domains highlighted on the 1-dimensional sequence of the spike protein as well as mutations defining Delta and Omicron BA.1 spike protein sequences. Domains are defined as such: head (13 to 1140), stalk (1141 to 1273), S1 (13 to 685), S2 (686 to 1140), N-terminal domain (NTD, 13 to 291), receptor binding domain (RBD, 331 to 528), receptor binding motif (RBM 437 to 508), subdomain 1 (SD1, 529 to 589), subdomain 2 (SD2, 590 to 675), furin cleavage site (FCS, 675 to 690), fusion peptide (FP, 817 to 834), fusion peptide proximal region (FPPR, 835 to 855), heptad repeat 1 (HR1, 910 to 984), central helix (CH, 985 to 1034), connecting domain (CD, 1035 to 1068), heptad repeat 2 (HR2, 1163 to 1210), transmembrane domain (TM, 1214 to 1234), and cytoplasmic tail (CT, 1235 to 1273). **(B)** Top and side views of the 1-RBD-up spike head highlighting domains defined in panel A. Domains are shown as space filling surface on chain A of the spike, chains B and C are represented with yellow ribbons. Glycan atoms are shown in light grey licorice. **(C)** Positions of Delta and Omicron BA.1 mutations on spike head structure. The Ancestral strain spike protein surface is shown in yellow ribbons (chain A) and space filling surfaces (chains B and C). Spike glycans are shown in light grey licorice. Mutations to the Ancestral strain corresponding to Delta and Omicron BA.1 spike sequences are shown as purple and blue beads, respectively. Mutations to the Ancestral strain shared between Delta and Omicron BA.1 spike sequences are shown in cyan beads.

Understanding spike biophysics can reveal adaptive pressures driving SARS-CoV-2 evolution.^36–38^ Following the initial sequencing of the SARS-CoV-2 genome in late 2019, and the subsequent spread of this virus around the world, in February 2020 a new clade (G) of the novel SARS-CoV-2 emerged, characterized by 4 mutations including a key single substitution from aspartic acid to glycine at the spike protein’s 614 position.^39^ By the end of March 2020, this SARS-CoV-2 “D614G variant” rapidly became the predominant circulating strain worldwide.^39^ Although the D614G mutation position is over 100 residues and ∼75 Å from the RBM’s center of mass in the closed state, initial epidemiological investigations revealed the D614G variant was between 4 and 9 times more infectious than the original novel 2019 strain, hereafter referred to as the Ancestral strain.^39^ Furthermore, several groups also noted D614G variant SARS-CoV-2 correlated with higher viral titers,^39–41^ higher S incorporation on the virion,^42–44^ and decreased premature S1-shedding^42^ on D614G viruses compared to the Ancestral strain. Rapidly, several groups sought to explain these observed differences through structural characterization of the D614G spike protein. While it is still unclear whether the D614G mutation results in an increase,^45,46^ decrease,^41^ or no change^42,43^ in ACE2 binding affinity, many groups identified that the D614G mutation correlates with increased populations of 1-up, 2-up, and 3-up RBDs relative to the Ancestral strain spike protein.^41,44,47,48^ Atomic scale structural inspection via high resolution Cryo-EM structures enabled predictions that D614G abolishes either a hydrogen bonding interaction to T859^45^ or salt bridging interaction to K854^44,45^ side chains. T859 and K854 are both located next to 614 on the neighboring chain’s S2 domain within the Fusion Peptide Proximal Region (FPPR, **Figure 1AB**). Zhang and Jackson et al. posited that the loss of D614’s interactions to the neighboring chain allow for increased conformational flexibility in this region potentially affording optimized interactions between S1 and S2 domains and stabilization of the spike trimer thus preventing premature S1 shedding.^42^ Zhang and Cai et al. showed that D614G substitution results in slight outward motion for the SD1 domain, exposing a larger volume within which the so-called “630loop” can arrange more tightly toward the core in a partial helix.^44^ Similarly, through molecular modeling and MD simulations, Dokainish and Sugita then showed that this same 630loop adopts a disordered structure in Ancestral strain spikes, but mutation to G at the 614 position allows the 630loop to become more ordered and stabilized.^49^ Interestingly, that Dokainish and Sugita^49^ and Yang et al^47^ probed the potential for pH regulation on spike conformational states by way of breaking the salt bridge to K854 through D614 protonation. Despite these, and many other, extensive investigations on D614G spike structural biology, the allosteric relationship between D614G mutation and RBD openness -- thus potentially increased probability for ACE2 binding, cell fusion, and infectivity – has not yet been clearly explained.

By the summer of 2020, the D614G variant was the predominant circulating strain of SARS-CoV-2.^39,50^ D614G’s increased infectivity and faster replication time then allowed for a dramatic expansion in genomic variation and emergence of many other variants of concern (VOCs), all incorporating the D614G spike mutation.^50^ A complete review of all strains is outside the scope of this work, however two strains of particular note include the Delta and Omicron BA.1, first detected in October 2020 and November 2021, respectively.^38,51,52^ The Delta spike glycoprotein is characterized by 9 mutations to the Ancestral spike sequence, **Figure 1**, including one mutation (T19I) which results in the loss of an N-linked glycan at position 17.^53,54^ Furthermore, early structural investigations revealed a refolding of several loops within Delta spike’s NTD, including significant rearrangement of the position of the N149 glycan,^26,53,54^ likely contributing to decreased immune recognition of Delta spikes.^54^ While no large scale structural rearrangements are seen in Delta RBDs or RBMs, two mutations within Delta’s RBM (L452R, T478K) are predicted to increase Delta’s binding affinity to ACE2^55,56^ and the P681R and D950N mutations are predicted to enhance FCS cleavage and membrane fusion, respectively.^57,58^ Similarly, the Omicron BA.1, hereafter referred to simply as “Omicron”, strain’s spike sequence is characterized by 34 mutations relative to the Ancestral spike sequence, **Figure 1**. These 34 mutations, including three deleted regions and an insertion, challenged COVID19 tracking efforts due to spike sequence dropout.^59–61^ Once the Omicron genome was sequenced and monitoring enabled, it was revealed that the Omicron SARS-CoV-2 variant was highly infectious, dwarfing infection and hospitalization rates from all earlier strains.^60–62^ Omicron’s high infectivity was hypothesized to be due to (1) the high number of sequence modifications, particularly in the RBD and NTD (15 and 8 mutations, respectively) leading to immunogenic escape,^33,63^ and (2) the high number of mutations in the RBM (10 mutations) potentially increasing binding affinity to ACE2.^64,65^ Additionally, Cryo-EM micrographs and epitope mapping reveals that Omicron RBDs are more likely to be in the up/open conformations relative to Ancestral strain spikes.^66^

Molecular dynamics (MD) simulations have proven valuable during the SARS-CoV-2 pandemic, particularly in revealing structure/function/dynamics relationships inherent to the spike glycoprotein and its otherwise “invisible” glycan motifs.^19–21,67^ Through conventional MD simulations, confirmed by biolayer interferometry (BLI) experiments, our group predicted the role of glycans at positions N165 and N234 in, like kick-stands on a bicycle, stabilizing RBDs in the up conformation.^19^ We have also used weighted ensemble (WE) enhanced sampling MD simulations to efficiently simulate the RBD opening conformational change.^21^ Our simulations revealed the key role of the N343 glycan in wedging the neighboring RBD into an open conformation by exploiting a hydrophobic stacking network.^21^ BLI experiments confirmed that loss of the glycan at this 343 position results in decreased ACE2 binding affinity likely due to an decreased number of up RBDs and exposed RBMs.^21^ Additionally, from these WE simulations we were able to characterize several key salt-bridge and hydrogen bonding interactions that are broken and/or formed during RBD opening, and we posited how mutations to, or nearby to, such interacting residues may impact RBD opening dynamics.^21^

The WE method enhances the sampling of barrier-crossing events by running multiple trajectories and periodically applying a resampling procedure based on a progress coordinate.^68,69^ In the resampling procedure, trajectories that make significant progress are duplicated, while those that do not are terminated. This approach is analogous to assigning more resources to a search team near a promising location. Importantly, each trajectory is assigned a statistical weight, and these weights are rigorously tracked to ensure that no bias is introduced in the dynamics.^68,69^ In this study, we used WE simulations with a minimal adaptive binning scheme^70^ to efficiently model RBD opening in the Ancestral, Delta, and Omicron spike glycoproteins. Our simulations reveal a key salt bridge (D614 to K854) that breaks before the RBD of the Ancestral strain can open. We also reveal an allosteric network that transmits information from the NTD to the opening RBD within the same chain and demonstrate how this network is strengthened by the D614G mutation. Additionally, we discuss biophysical factors such as the flexibility of the 630loop and relative RBD motion in the Ancestral, Delta, and Omicron spikes.

## Results and Discussion

We performed equilibrium WE simulations of the Ancestral, Delta, and Omicron spike proteins similarly to those described by Sztain et al (2021)^21^ wherein we defined RBD opening according to the following two-dimensional (2D) progress coordinate: (1) the distance between the center of mass of the RBD β-sheet C_α_ atoms to the center of mass of the spike’s central helix C_α_ atoms, referred to as the RBD-core distance (**Figure 2A**), and (2) the root mean square deviation (RMSD) between the RBD β-sheet C_α_ atoms in each simulation frame and the RBD β-sheet C_α_ atoms seen in PDB ID 6VSB, referred to as RMSD to 6VSB.^9^ PDB ID 6VSB was the first deposited SARS-CoV-2 spike ectodomain structure and is a well-known pre-fusion stabilized open RBD spike structure that served as the basis for several early SARS-CoV-2 mRNA vaccines,^9^ making it an ideal reference for guiding our RBD opening simulations.

**Figure 2.**
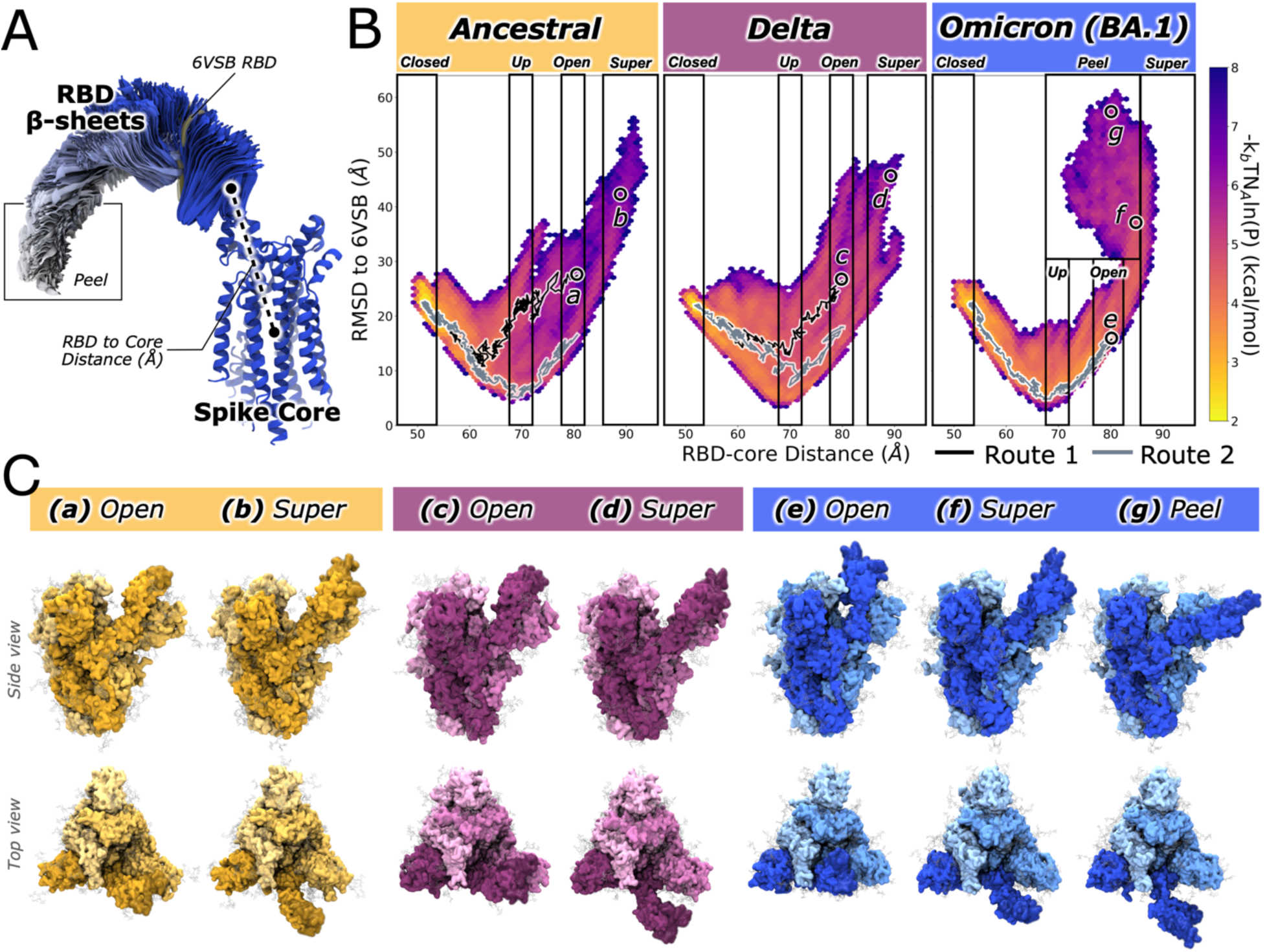
Conformational landscapes for Ancestral, Delta, and Omicron BA.1 spike openings. **(A)** Progress coordinates used in this work to encourage RBD opening via WE simulations. To highlight Omicron’s RBD Peel conformation, Omicron opening trajectories were used to develop this image. The spike core is defined by α-helical bundle within the center of the spike (747 to 785, 945 to 1035) is represented as dark (chain A) and lighter (chains B and C) blue ribbons. The center of mass of C_α_ atoms within the spike RBD β-sheets is used to define the RBD position relative to the center of mass of C_α_ atoms within the spike core. RBD β-sheets (residues 353-358, 375-381, 394-403, 430-438, 507-516) are colored according to their conformational state: closed to up conformations in dark blue, open conformations in light blue, super-open and peel conformations in white. RMSD of RBD β-sheet C_α_ atoms between simulation frames and an early standard 1up spike structure (6VSB) was also used to drive RBD opening. As such, the position of 6VSB’s RBD β-sheets is highlighted. **(B)** Histograms for Ancestral, Delta, and Omicron spikes revealing the relative probability (kcal/mol) for RBD conformations along opening pathways as a function of the two-dimensional progress coordinate. Regions of conformational landscapes corresponding to our defined closed, up, open, super (super-open), and peel states are highlighted. Representative pathways of routes 1 and 2 (black and grey respectively) are overlaid on top of the conformational landscapes. **(C)** Representative conformations of the open, super, and peel states for the Ancestral **(a,b)**, Delta **(c,d)**, and Omicron BA.1 **(e,f,g)** spikes. Positions of these conformations within the entire landscape are highlighted in panel B.

Our WE simulations employ the minimal adaptive binning (MAB) scheme,^70^ which dynamically identifies bottlenecks and adjusts bin positions during the simulation. To ensure comparability between Delta and Omicron RBD opening results performed in this study, we thus also conducted Ancestral RBD opening simulations using this MAB scheme.

### Ancestral, Delta, and Omicron spikes traverse distinct RBD opening landscapes

Upon completion of WE simulations for Ancestral, Delta, and Omicron BA.1 spike RBD opening we immediately observed that the three spikes traverse different regions of phase space as defined by the 2D progress coordinates, **Figure 2B**. During the RBD opening process, as RBD’s open and RBD-core distance increases, the RMSD to 6VSB decreases. All spikes reach a minimum of ∼5 Å in RMSD to 6VSB corresponding to an RBD-core distance of ∼70 Å in the up state. All RBDs then continue to open further beyond 6VSB’s position, thus as the RBD-core distance increases beyond ∼70 Å the RMSD to 6VSB also increases as simulation frames again become different from the 6VSB state and spikes move into the open and super-open states. To estimate the relative rates of RBD opening for each spike variant, we stitched complete successful trajectories moving from closed to up and open target states and calculated the total time of each trajectory. These results, **Table 1** and **Figure S1**, suggest that Delta and Omicron spikes rapidly move into up and open conformations, while Ancestral spikes lag by an average of ∼20-30 ns when moving to up and open states.

**Table 1.** Mean relative molecular times calculated from closed to up and open states, all times listed in ns.

| Spike | Closed to Up | Closed to Open |
| --- | --- | --- |
| Ancestral | $40.59 \pm 3.17$ | $45.45 \pm 1.69$ |
| Delta | $11.68 \pm 2.78$ | $16.53 \pm 2.73$ |
| Omicron BA.1 | $17.70 \pm 6.54$ | $20.17 \pm 4.55$ |

Despite the above-described overarching similarities within the 2D reaction coordinate, there are clear distinctions in the reaction profile silhouette between the three strains. While Ancestral and Delta spikes demonstrate higher variability in RMSD to 6VSB values (5 to 30 Å) within an RBD-core distance range of 55-70 Å, the Omicron spike traverses a more direct path within this region, occupying a tighter RMSD to 6VSB range between 5 and 20 Å, **Figure 2B**. Furthermore, while we observe that all spikes visit the up (RBD-core distances 68-72 Å), open (RBD-core distances 78-82 Å), and super-open (RBD-core distances >85 Å) states (**Figure 2C**) with relatively high probability, the Omicron RBD moves into the “super-open” state but then adopts conformations with decreasing RBD-core distance values, i.e., the Omicron spike opening profile almost appears to double back on itself. Visualizing these “doubled-back” Omicron conformations we see that the Omicron RBD is opening so far that, like peeling an orange, the RBD’s center of mass begins to move down towards to the spike’s central plain, thus decreasing the RBD-core distance. As such, hereafter we will refer to this region of conformational space as the Peel state.

### Clustered opening pathways reveal diversity in Ancestral and Delta pathways but similarity in Omicron BA.1 pathways

While large ensembles of pathways, as shown in **Figure 2B**, are invaluable for describing the nuanced variability in RBD opening for Ancestral, Delta, and Omicron spike proteins, these data sets can be cumbersome to analyze (complete trajectories from Ancestral, Delta, and Omicron BA.1, with water molecules and ions removed, are ∼660GB, ∼350GB, and ∼320TB in total size, respectively, **Table S1**). To mitigate these challenges, we applied the Linguistics Pathway Analysis of Trajectories with Hierarchical Clustering (LPATH) method to cluster all successful RBD opening pathways into distinct routes (see Supporting Information Methods Section 1.2.2). From these results we identified two pathway routes for each of the Ancestral, Delta, and Omicron spike proteins, **Figure 2B** (black and grey overlayed trends) and **Figure S2**.

As can be seen in **Figure 2B**, the Ancestral spike’s routes 1 and 2 are very distinct. As the RBD opens (RBD-core distance increases) route 2 first directly minimizes RMSD relative to the 1-RBD-up 6VSB structure, however route 1 initially minimizes RMSD to 6VSB but diverges early on from route 2 at RBD-core distance ∼60Å and begins to increase RMSD to 6VSB before completing the RBD opening. Similarly, the Delta spike’s routes 1 and 2 are distinct in that route 2 follows the lower profile monotonically minimizing RMSD to 6VSB, whereas route 1 diverges from route 2 at RBD-core distance ∼70Å by increasing RMSD to 6VSB. These results suggest Ancestral and Delta spike proteins explore two types of opening pathways: one wherein their RBD position is relatively more similar to 6VSB (route 2 in Ancestral and Delta spikes) and one wherein their RBD positions are distinct from 6VSB (route 1 in Ancestral and Delta spikes). However, the two routes extracted from Omicron RBD opening trajectories are very similar to one another in progress coordinate space: they both minimize RMSD to 6VSB while the RBD is opening. As such, there are likely other conformational changes in Omicron RBD opening which are not described along the 2D progress coordinate space but for which LPATH analysis is sensitive enough to differentiate, i.e., Omicron’s routes 1 and 2 may be distinct along an axis not immediately discernable via the progress coordinate landscape. Interestingly, these results support our hypothesis gleaned from the full landscape results: Omicron RBD opening follows a more direct set of routes with high similarity to one another in early RBD opening stages, whereas Ancestral and Delta RBD opening routes demonstrate lower similarity, higher variability, in early stages of RBD opening. Interestingly, we can also see that routes 1 and 2 for Ancestral and Delta spike proteins both diverge, as described, from one another but that divergence occurs in different regions of progress coordinate space. For the Ancestral spike, route divergence occurs around an RBD-core distance of ∼60 Å. For the Delta spike, clear divergence between routes occurs once the RBD has already reached the up state around an RBD-core distance of 70 Å. The implications of Ancestral RBD opening divergence at 60 Å versus Delta divergence ∼70 Å and Omicron’s apparent lack of divergence is not yet clear.

### The N2R and R2N flexible linkers unite all major S1 domains

Gobeil and Henderson et al identified conformational correlations between RBD openness and a linker peptide which they designated the NTD-2-RBD (N2R), residues 293 to 330.^71^ As can be seen in **Figure 3** (pink ribbon), beginning approximately at position 293, the N2R linker runs from the base of the NTD, serves as a β-strand connected to the SD2 (K310 to T315), forms a short loop (Ancestral) or β-strand sheetlet (Delta and Omicron) (T315 to P322), serves as a β-strand within the SD1 (P322 to N331), and finally passes into the RBD (N331) as a flexible loop around a previously reported transient pocket.^72^ The RBD itself is then largely well folded into a bundle of α-helices surrounding a central β-sheet and fortified by via several disulfide bonds (C336-C361, C379-C432, C391-C525), including one within the RBM (C480-C488). The one-dimensional spike sequence representation, however, belies the fact that the S1 folds back onto itself: The loop exiting from the RBD (P527-C538, lilac ribbon, **Figure 3**) runs adjacent to the SD1 and directly connects to another β-strand (C590 to G594) – the anti-parallel β-strand paired with N2R’s T315 to P322 peptide – via a disulfide bond (C538-C590), and then wedges between the base of the NTD and the SD2 via another β-strand (G594 to T602) within the SD2’s central β-sheet. Herein, we will refer to this second linker that connects the RBD back to the base of the NTD via a disulfide bond as the R2N linker (residues 526 to 538, C538-C590, and 590 to 602), which runs parallel to and engages significantly with the N2R (residues 293 to 330). The N2R and R2N both pass over the SD1 to the RBD and ultimately staple to SD2’s β-sheet. In keeping with the trend in previous spike structural biology investigations likening the spike domains to joints -- like the ankle, knee, and hip^73^ -- one could liken the N2R and R2N linkers as “tendons” extending through the spike elbow (NTD to SD2 to SD1) and wrist (SD2 to SD1 to RBD) joints. Gobeil and Henderson et al show clearly that, for Omicron strain spikes, N2R linker position is highly correlated with RBD openness.^71^

**Figure 3:**
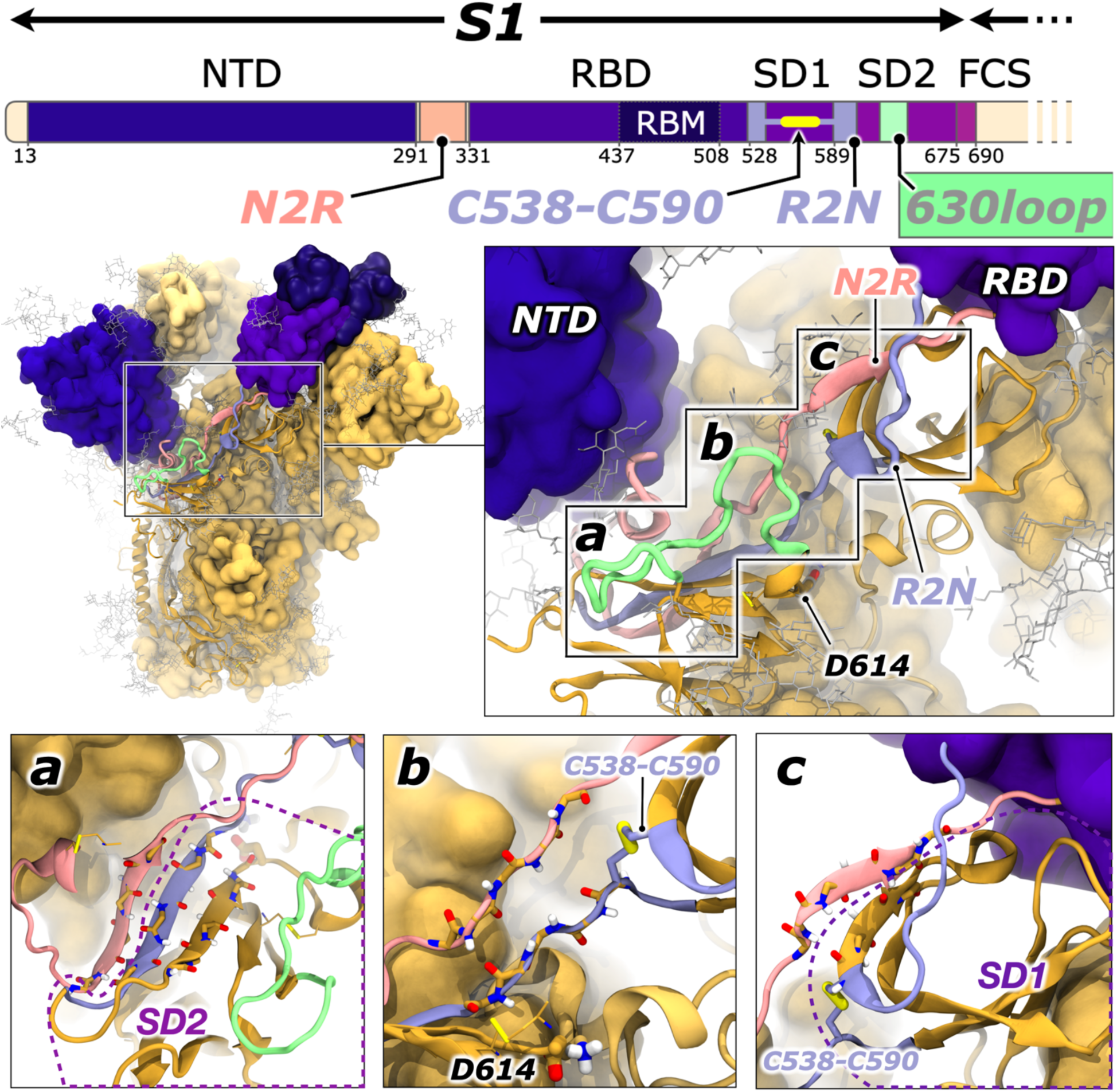
Details of the N2R and R2N linkers and the 630loop. A one-dimensional sequence of the spike S1 shows how the N2R linker (pink ribbons) connects the NTD directly to the RBD. The R2N linker (lilac ribbons) is revealed when considering the covalent link via the C538 and C590 disulfide bond. The 630loop (green ribbons) has been described by others to correlate in motion and stability with RBD opening. Panels a, b, and c depict the backbone atoms in licorice (carbon, oxygen, nitrogen, and hydrogen in yellow, red, blue, and white, respectively) to highlight the regions in which the N2R and R2N interact with beta-sheets within the SD2 (panel a) and SD1 (panel c) or pass through the 614 proximal region. For clarity, the NTD is not displayed in panel a, and the 630loop is not displayed in panel b.

Gobeil et al^71^ and Manrique et al^74^ have also identified loops within the N2R as likely important in facilitating spike RBD conformational changes. Furthermore, despite the fact its position is far from the RBM (∼75 Å), several groups have reported the D614G mutation correlated significantly to increased infectivity of SARS-CoV-2.^39–49,75^ Therefore, the relationship between position 614, ACE2 binding, and SARS-CoV-2 infectivity suggests long-range allosteric communication. While not located in the N2R or R2N, the 614 position is directly proximal to a β-sheetlet within the N2R and R2N (T315 to P322 and C590 to G594, respectively). Thus, considering the likelihood of N2R and R2N linkers facilitating allostery within the S1, and the proximity of these linkers to the 614 position, we then sought to characterize correlated motions and interaction differences within these linkers. Furthermore, Zhang et al^44^ and Dokainish and Sugita^49^ have noted distinct structural and dynamical changes for the so-called 630loop within the SD2: this loop extends off the β-turn where 614 sits, moves around the SD2 and reconnects to the beta-sheet. Dokainish and Sugita^49^ showed that not only was the position of the loop correlated to RBD opening, but also that the D614G mutation, or protonation of D614, altered stability of the 630loop allowing tighter packing toward the spike core.^49^ Zhang et al also showed the 630loop becomes more ordered in Omicron and later spikes.^76^

### Enhanced allosteric communication within Delta and Omicron spikes

To identify and evaluate the strength of allosteric communication networks between linkers and domains during RBD opening, we performed dynamical network analyses using the Weighted Implementation of Suboptimal Paths (WISP).^77^ Using graph theory, we constructed residue-based networks using cross-correlation analysis where nodes and edges are weighted by their strength of correlated motions^77^ (for complete methodological description see Supporting Information Methods Section 1.2.5). This approach therefore not only predicts allosteric networks but also estimates how the strength of correlation within these networks can change as a function of the sequence mutations and structural rearrangements as seen in variant spikes. For all spike RBD opening routes, we see differences in the predicted allosteric networks within the chain undergoing RBD opening as a function of variant, **Figure 4**, as well as differences within the non-opening chains and between chains, **Figure S3**. Herein, we will describe differences seen within the RBD-opening chains for the routes with highest probability (as taken by the relative weight per route trajectory selected via LPATH). As shown in **Figure 4B**, for the Delta spikes, allosteric communication is relayed between the NTD and RBD via the R2N linker (lilac spheres and rods), and for the Omicron (**Figure 4C**) spike allosteric communication is relayed via the N2R (pink spheres and rods) and the R2N linkers. However, for the Ancestral spike, the allosteric pathway connecting the NTD to the RBD relies primarily on the N2R (**Figure 4A**), with very little communication through the R2N. Interestingly, in the Ancestral spike, only one node within the R2N is activated in the less probable route 2 (**Figure S3A**) where a strong degree of correlated motion is observed between residues N317 and F592.

**Figure 4.**
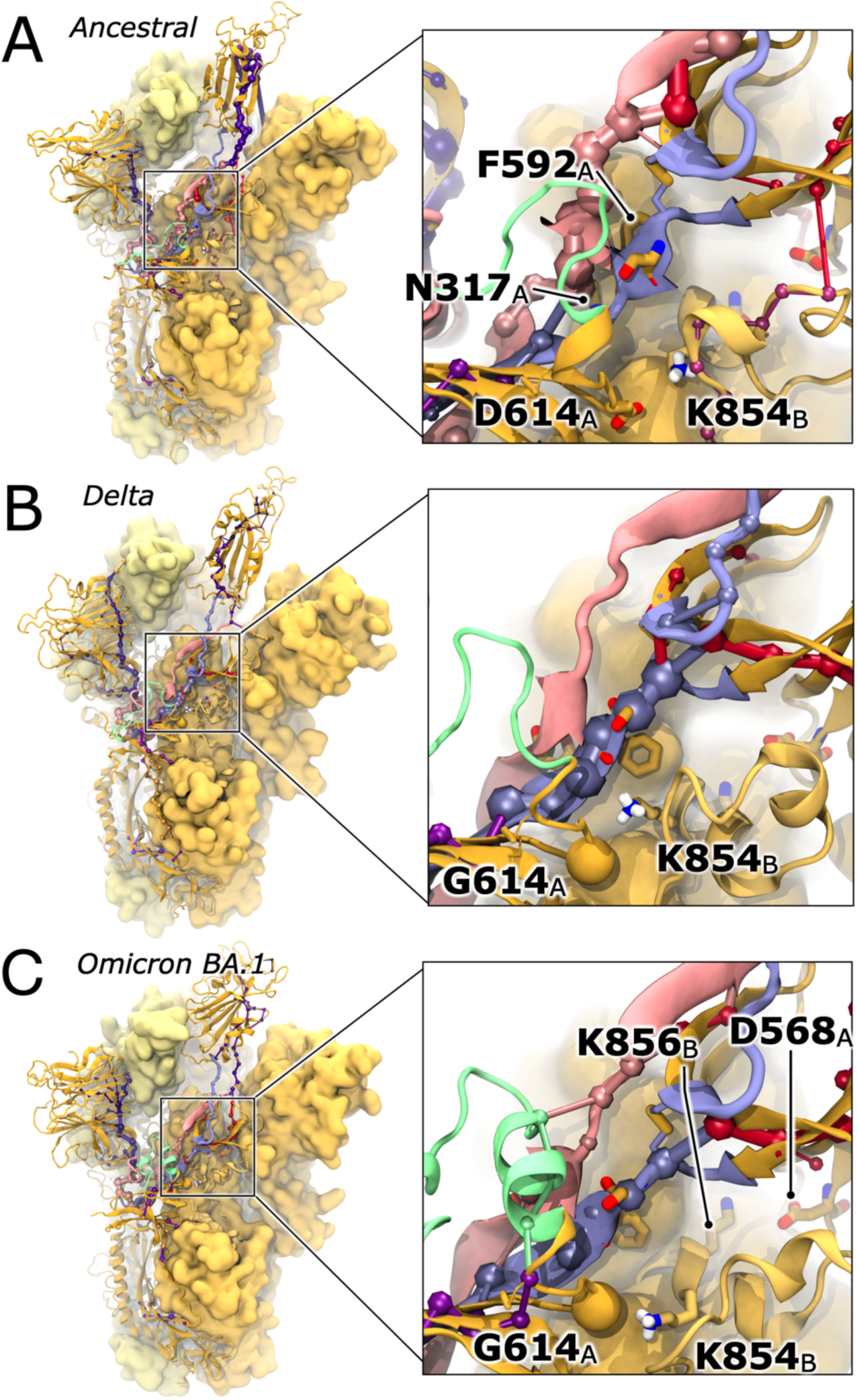
Dynamical networks reveal new lane of communication in Delta and Omicron Spike proteins. Dynamical networks calculated for **(A)** Ancestral, **(B)** Delta, **(C)** Omicron BA.1 spike proteins from the most probable opening pathways. In all panels, the spike is represented in yellow ribbons (chain A) and medium- and light-yellow surfaces (chain B and C, respectively). The FP and FPPR of chain B are represented as a yellow ribbon to reveal positions of K854. Correlated motion networks are shown as spheres and edges and baseline networks are depicted according to which domain or linker the node belongs, for Ancestral, Delta, and Omicron spikes, respectively. Amino acids of interest (residues 614, 317, 592, and 854) are represented in licorice with yellow carbon atoms.

Conversely, in Delta spike’s most probable RBD opening route (2), the R2N facilitates the allosteric communication between the NTD and RBD, **Figure 4B**, with very little contribution from the N2R. Delta’s less probable, but still likely, route 1 does reveal significant communication within the N2R, **Figure S3B.** Thus, the precise RBD to NTD communication network may vary depending on the pathway traversed by the RBD during opening, but for Delta spikes the R2N is the primary facilitator of this allosteric information. Interestingly, in both of Omicron’s RBD opening routes, we observe clear and robust communication through both N2R and R2N linkers, **Figure 4C**. In fact, communication networks for both Omicron routes show only very minor differences. This is consistent with earlier descriptions of the complete conformational landscapes collected from Omicron simulations, wherein the Omicron RBD opening follows a "tighter" probability profile, transitioning more directly from closed to partially up and open before exploring broader conformational space in the super-open and peel states (**Figure 2A**). Omicron’s nearly identical allosteric networks for both pathway classes suggests that early stages of Omicron RBD opening follow a more consistent route compared to Ancestral and Delta, where the RBD explores a broader range of positions relative to the 6VSB structure (RMSD to 6VSB).

Contrary to Delta and Omicron, allosteric networks in the Ancestral spike relies largely on correlation through the N2R only: 34.3% and 38.4% of total weights for nodes within Delta and Omicron allosteric networks pass through the R2N, respectively, compared to 11.4% through the Ancestral spike’s R2N. Additionally, Delta and Omicron spikes exhibit more balanced communication between N2R and R2N linkers compared to the Ancestral spike: node weights for the N2R and R2N within Ancestral spike were 87.8% versus 11.4%, compared to 64.8% and 34.3% for Delta, and 58.7% and 38.4% for Omicron. Thus, much like opening new lanes on a highway to alleviate congestion, significant degrees of communication flows through both the R2N in Delta and Omicron spikes during RBD opening, likely allowing for more direct, concerted, and attenuated communication pathways between NTDs and RBDs. Additionally, the Omicron spike’s 630loop is directly connected to the RBD-to-NTD allosteric network, whereas this same region is largely disconnected from this allosteric network in Ancestral and Delta RBD opening pathways, **Figure 4** and **Figure S3-4**. As mentioned, several groups have posited a correlation between the 630loop motion and stability to RBD opening.^44,49^ Here, we see that Omicron’s 630loop is very well ordered with two additional α-helical turns relative to the Ancestral spike. Additionally, we see that during RBD opening, the 630loop stays far more compacted towards the spike core in Omicron spikes than in Delta or Ancestral spikes and that compaction towards the core only increases (decreasing distance from core) during RBD opening on the chain that is opening, **Figure S5.**

### The D614 to K854 salt bridge dampens S1 allostery and must break before the Ancestral spike’s RBD can open

We believe that communication along the R2N in the Ancestral spike is likely hindered due to steric congestion caused by a neighboring salt bridging interaction between D614 and K854. As mentioned previously, the N2R and R2N linkers pass directly “behind” the 614 position. We sought to characterize the degree to which this salt bridge and the 614 position may impact allostery through the 614 proximal region. Cryo-EM resolution of the D614G spike structure, and comparisons of this structure to Ancestral strain spike structures, reveal potential interactions between D614 and two residues, T859 and K854, within the neighboring chain’s FPPR.^45^ These interactions have been posited as likely stabilizing the RBD while it is opening.^44^ Considering the potential of the D614G mutation to alter spike allostery, we carefully probed the region around the 614 position to uncover any relationship to RBD dynamics.

Upon first observing our stitched RBD opening simulations, we quickly identified that in the closed Ancestral spike, D614 engages the neighboring chain’s K854 in a tight salt-bridge, **Figure 5AB**. As can be seen in **Figure S6,S14B**, for the Ancestral spike, the D614-K854 salt bridge is tight in the closed state (RBD-core distance < 55 Å) but breaks in initial stages of transition from closed to up states (RBD-core distances 55 – 65 Å). The D614-K854 salt bridge then reforms in later stages of this transition (RBD-core distances 65 – 68 Å) and is again tightly engaged again when the RBD is in the up state (RBD-core distances 68 – 72 Å). Consistent with our results, several Cryo-EM spike structures in up and open conformations show the D614-K854 salt bridge is either tightly engaged,^44,78^ or could be engaged if the FPPR were resolved, but our simulation results can capture transition state conformations wherein the salt bridge breaks and reforms in the 1up RBD state. Beyond the Ancestral spike’s up state, as a function of the 2D conformational landscape, the D614-K854 salt bridge occupies regions of tight, light, or no engagement depending on RBD position. Furthermore, in all successful pathway trajectories for Ancestral spike opening, we noted that the D614-K854 salt bridge breaks before the RBD can make significant progress along the opening pathway. As described earlier, the Ancestral spike spends significant amounts of time (44.5+/-3.1% of frames, **Figure S14A**) per stitched opening trajectory in the closed conformation (RBD-core distance < 55Å). Most of the Ancestral spike’s closed frames show a tightly engaged salt bridge (93.1+/-0.0% of all closed frames from stitched opening trajectories have D614-K854 salt bridge distance < 3Å), **Figure S6**. In fact, most Ancestral opening pathways spend ∼17ns of molecular time prior to finding the first broken D614-K854 salt bridge distance (D614-K854 distance > 4Å), and only another ∼5ns of molecular time (total ∼22ns) before the salt bridge sustains a broken conformation (more frequent than every 4^th^ frame with a D614-K854 distance > 4Å). At ∼24.5ns, with the D614-K854 salt bridge in the sustained-broken conformation, the Ancestral spike moves out of the closed state into the transitional state heading towards the up state (RBD-core distance > 55 Å, **Figure 5B Transition** and **Figure S14AB, S6)**. Around 30ns of molecular time, D614 and K854 move even further apart (> 5Å) and concomitantly the Ancestral RBD progresses linearly (∼0.8Å/ns) towards the up state (RBD-core distances 55 – 62 Å, **Figure S14AB, S6**). At ∼36ns, the D614-K854 salt bridge reforms (< 4Å), slowing down but not halting RBD opening at ∼0.5Å/ns (RBD-core distances 62 – 68 Å, **Figure 5B Up/Open** and **Figure S14AB, S6**). As described above, in the RBD up state (RBD-core distances 68 – 72 Å, **Figure S6**) the D614-K854 salt bridge reforms, again as seen in several Cryo-EM structures.^44,78^ Beyond the up state, there are several pockets of conformational space in which the D614-K854 salt-bridge is more or less engaged, suggesting RBD position in the Ancestral spike is highly correlated with the strength of this interaction.

**Figure 5:**
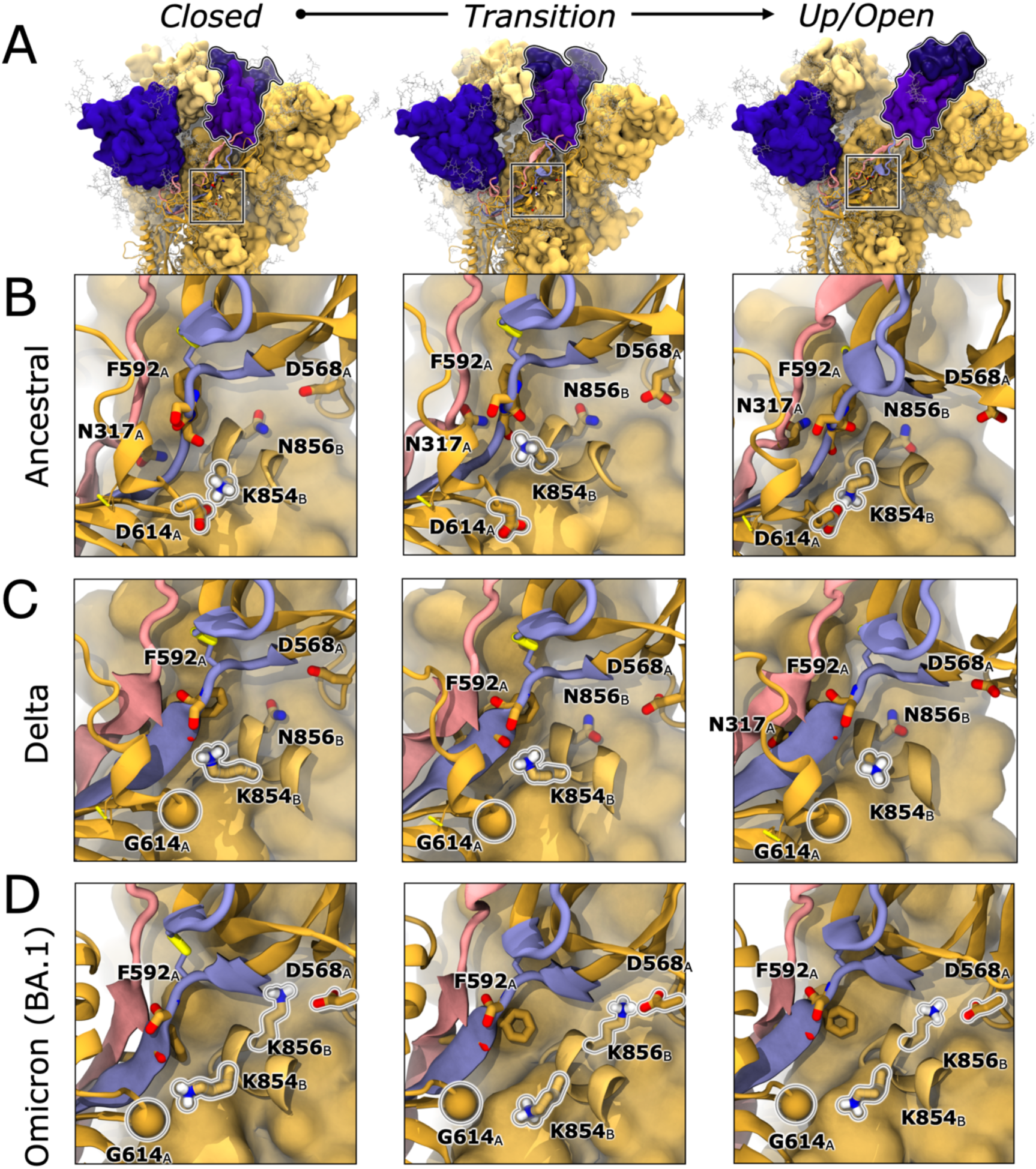
RBD opening progress as a function of salt bridging interaction between D614 and K854. In all panels, the spike is represented in dark yellow ribbons (chain A) and medium- and light-yellow surfaces (chain B and C, respectively). The FPPR of chain B is represented as a medium yellow ribbon to reveal positions of K854 and N/K856. The N2R (residues 293 to 330) and R2N (residues 526 to 538, C538-C590, and 590 to 6023) are highlighted in pink and lilac ribbons respectively. Amino acids of interest (residues 614, 317, 592, 854, 856, and 568) are represented in licorice with dark and light yellow carbon atoms indicating chain A or B, respectively. Disulfide bonds are highlighted with thick yellow licorice. The G614 position in Delta and Omicron spikes is denoted with a yellow sphere. **(A)** Frames from Ancestral spike opening trajectories were selected from closed, transition, and open states to highlight RBD opening progress correlating with the zoom-in images on the 614 proximal region for the **(B)** Ancestral, **(C)** Delta, and **(D)** Omicron BA.1 spike proteins. In the closed and open states for the Ancestral spike, the D614 to K854 salt bridge is tightly formed. Before the RBD can begin to open the D614-K854 salt bridge breaks and remains broken until the RBD enters the up conformation (supporting information **Figure S6,S14B**). For Delta and Omicron spikes, the D614 to K854 salt bridge is ablated and thus there is no reliance on that interaction breaking before RBD opening. In the Omicron spike, a new salt bridge is established between K856 and D568, and this salt bridge remains intact throughout RBD opening.

Conversely, the D614G mutation, as seen in Delta and Omicron spikes, ablates the salt bridge to K854. As such, for Delta and Omicron RBD opening simulations we see no correlation between G614-K854 distance and RBD-core distance or molecular opening times, **Figure 5CD** and **Figure S6,S14AB**. Additionally, as mentioned, Delta and Omicron spikes spend very little time in the closed state, (11.4+/-2.9%, 17.3+/-5.2% of RBD opening frames with < 55 Å RBD-core distance, respectively) and instead begin to open quickly at ∼1.3 Å/ns and ∼1.2 Å/ns respectively **(Figure S14AB, S6)**.

Following our characterization of the D614-K854 salt-bridge, we also probed neighboring interactions around 614’s position particularly any interactions with the N2R and R2N linkers. We saw that the Ancestral spike’s D614-K854 salt bridge breaking event was concomitant with a backbone twisting (as identified from φ/ψ distributions) observed for R2N-linker residues P589 to G594 (PCSFGG, **Figure S7,14E-H**), particularly S591 and F592, which is not observed for Delta or Omicron RBD opening pathways. Furthermore, in the Ancestral spike, we see an increase in contacts between N2R-linker residue N317’s side chain and R2N-linker residue F592’s backbone after the φ/ψ switch (**Figure S8,14D**) where this contact is not seen in Delta and Omicron spike proteins. Interestingly, during the RBD transition into the up-state, after the D614-K854 salt bridge reforms, backbone φ/ψ distributions do not revert to their conformation in the closed state, **Figure S7,14E-H**. Considering this backbone twist was only seen for Ancestral spikes, and its correlation with D614-K854 salt bridge breaking, we hypothesized that local steric congestion could be impacting R2N dynamics. Thus, we calculated the distance between the SD2 and R2N (particularly residues 590-594, **Figure S10**), the R2N (residues 590-594) and the neighboring chain’s FPPR (**Figure S11**), the SD2 and the neighboring chain’s FPPR (**Figure S12**), and the area between these three domains (**Figure S13**). For the Ancestral spike, we see that R2N residues 590-594 are closer to the neighboring chain’s FPPR by ∼1-4Å than in Delta and Omicron spikes in the opening protomer. We also see that the area between the SD2, R2N, and neighboring FPPR is ∼10-20 Å^2^ closer than that seen in Delta and Omicron spikes. Moreover, the area between the SD2, R2N, and the FPPR increases during RBD opening in a profile similar to that seen for the breaking of the D614-K854 salt-bridge, **Figure S14J-L**. These results suggest the D614-K854 salt bridge induces local steric congestion within the D614 proximal region around, and likely impeding optimal allosteric communication within, the R2N linker.

Considering this relationship between the D614-K854 salt bridge contact profile during RBD opening (**Figure S6,14B**), R2N’s observed backbone twist particularly at positions S591 and F592 (**Figure S7,14E-H**), significant contacts between N2R-linker residue 317 and R2N-linker residue 592 **(Figure S8,14D)**, and the decreased SD2-FPPR-R2N area for Ancestral spikes relative to Delta and Omicron (**Figure S10-13)**, we believe the strong D614-K854 salt-bridge in the Ancestral spikes disrupts R2N facilitated allosteric communication and instead requires all communication to pass only through the N2R-linker. Breaking of this salt-bridge via the D614G mutation, we believe, alleviates frustration in the 614-proximal region, and opens communication along the R2N as seen for Delta and Omicron spikes. What’s more, as described above, the Ancestral RBD does not begin opening until the D614-K854 salt-bridge breaks. As a result, when considering all stitched opening pathways, the Ancestral spike spends ∼44.5+/-3.1 % of frames in the closed state and 92.7+/-0.0% of that time is spent iterating until a sustained broken salt-bridge conformation (>4Å, more frequent than every 4^th^ frame) is found allowing the RBD to begin to emerge from the down state (RBD-core distance > 55Å). Conversely, Delta and Omicron spikes spend only 11.4+/-2.9% and 17.3+5.2% of stitched opening frames, respectively, in the closed state before their RBDs begin to emerge and we see no relationship between G614-K854 distance and RBD opening, **Figure 5** and **Figure S6**. Thus, we believe ablation of the salt-bridge by D614G opens a lane of allosteric communication along the R2N allowing for enhanced allostery by concerted correlated motions contributing to faster RBD opening for D614G spike variants, consistent with our relative molecular opening times **(Table 1, Figure S14A)**. Our results are consistent with those from Manrique et al wherein they performed conventional MD simulations of spikes in the closed state. Their results reveal several residues within the N2R have high betweenness along allosteric communication networks, indicating their importance.^74^ Our work builds on their results as we have confirmed the N2R allosteric network, even in the presence of N- and O-linked glycans, across several spike variants, and support that such communication networks are vital for RBD opening. Additionally, several groups have also shown the D614G mutation results in an increase in flexibility in the surrounding region, likely leading to increased RBD flexibility.^40–42,44,45,47–49^ Taken together, these results strongly suggest that D614G increases RBD opening rates by alleviating conformational frustration around the R2N in the 614 proximal region, thereby affording concerted and consistent NTD to RBD allostery via both N2R and R2N linkers.

Additionally, upon visualizing Omicron RBD opening trajectories we also observed the establishment of a new salt bridge between FPPR residue K856 (result of N856K mutation) and the neighboring chain’s D568, **Figure 5D**. We calculated the strength of this salt bridge during Omicron RBD opening and see that it is tightly engaged throughout all regions of the 2D progress coordinate space, **Figure S9,14I**. Due to the presence of the bulky, polar, yet uncharged N residue at the 856 position, Ancestral and Delta spikes demonstrate short N856-D568 distances, but those short distances do not reflect tight engagement and are not correlated to RBD-core distance. The 856 position is within the FPPR and near K854. Our results suggest the K856-D568 salt bridge may allow for re-stabilization of the FPPR via re-introduction of the salt bridge while still affording necessary flexibility around, and allosteric communication through, both N2R and R2N linkers.

### HDXMS confirms D614G altered dynamics around spike allosteric networks

Considering the contact and communication changes observed in N2R and R2N linkers around position 614, we sought to experimentally probe dynamics in this region via Hydrogen/Deuterium Exchange Mass Spectrometry (HDXMS) from SARS-CoV-2 spikes embedded on virus-like particles (VLPs) for Ancestral, D614G, and Omicron BA.1 strain spikes. (Note: each spike represents the circulating genetic sequence for the given variant of concern, i.e., recombinant spikes incorporating proline substitutions were not used.) We identified coverage of two peptides near D614G and N2R and R2N-linkers. Peptide A (**Figure 6A**), residues 300-311, contains a short α-helix leading to a flexible loop located at the base of the NTD. Peptide B (**Figure 6B**), residues 842-850, is within the fusion peptide proximal region (FPPR, residues 834-855), is near to position 854, and includes a loop and α-helix. Ancestral and Omicron spikes exhibit a similar degree of deuterium uptake for Peptides A and B, while D614G spikes on SARS-CoV-2 VLPs show dramatically decreased deuterium uptake, especially for Peptide B and at early time points.

**Figure 6:**
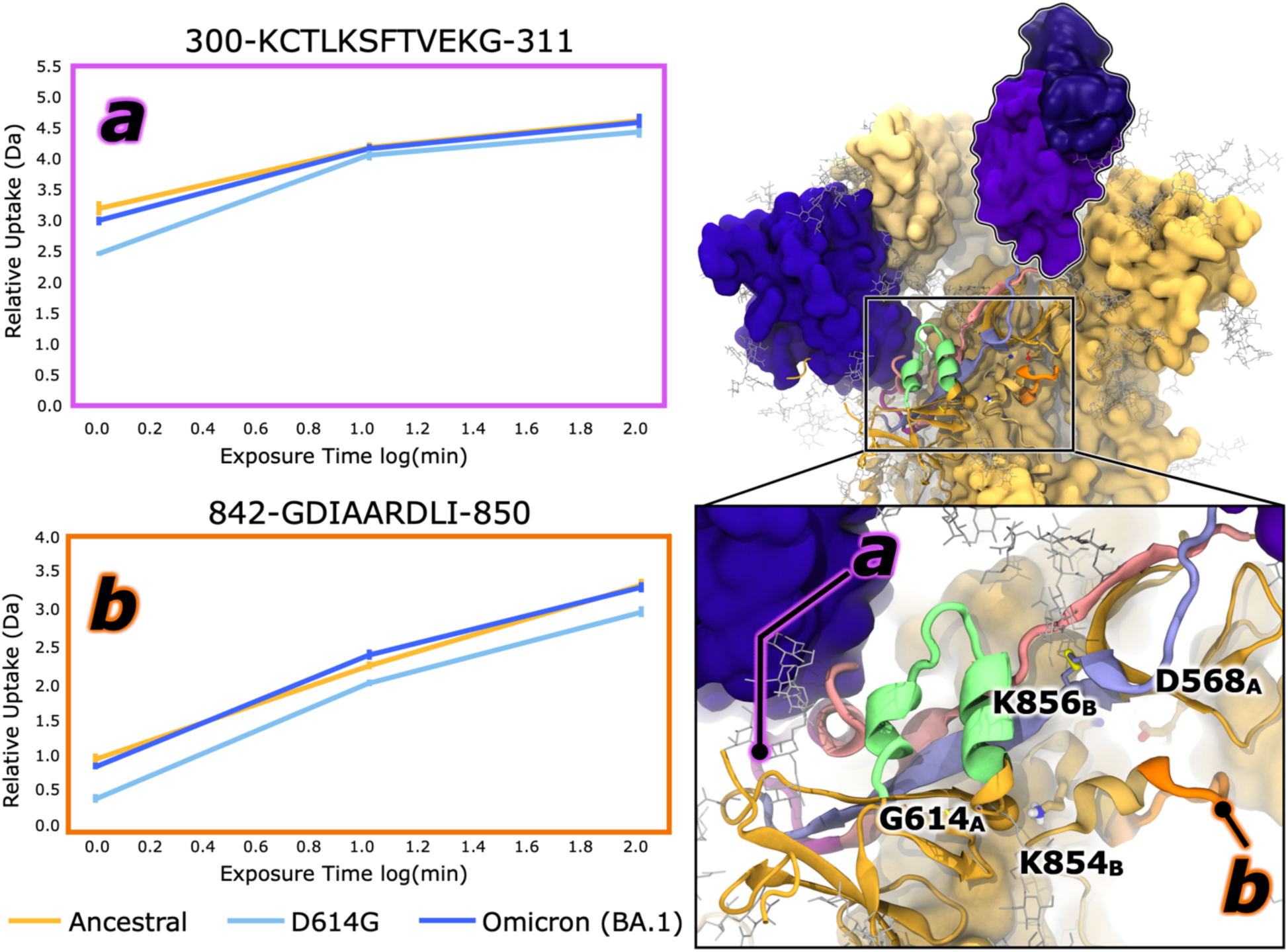
HDXMS results confirm structural and dynamic changes in 614 proximal region. Omicron BA.1 spike structure (right) is used to denote positions of peptides A and B relative to the NTD, RBD, FPPR, tendon 1 and tendon 2. Spike structure is shown in yellow ribbons (chain A) and medium- and light-yellow surfaces (chains B and C, respectively). As done in Figures 2 and 3, the N2R, R2N, and 603loops are highlighted in pink, lilac, and green ribbons, respectively. Amino acids of interest (residues 854, 856, 568) are shown in licorice with dark or medium yellow carbon atoms indicating chain A and B, respectively. Disulfide bonds are highlighted with thick yellow licorice denoting sulfur atom positions. Magenta and orange ribbons highlight the position of peptides A and B. Relative uptake plots, depicting deuterium uptake of peptides A and B, expressed in Daltons as a function of log(mins), are shown (left). Ancestral, D614G, and Omicron BA.1 spike data is represented in yellow, cyan, and blue trends, respectively.

Our HDXMS results support that the D614G mutation impacts NTD and FPPR dynamics on the minutes timescale, which is consistent with the experimentally observed rate for RBD opening, which is on the order of seconds.^79^ Considering our simulation results demonstrating the loss of the salt bridge contact and enhancement of communication along the R2N, we believe the base of the NTD and the FPPR loop compensate via rigidification, thus explaining loss of deuterium uptake in these regions in D614G compared to Ancestral spike proteins. However, with re-establishment of a salt bridge in the region with Omicron’s N856K mutation, Peptides A and B can return to similar degrees of flexibility as seen in the Ancestral strain. Although a comprehensive characterization of backbone hydrogen bonding in this region is outside the scope of this current work, we believe these HDXMS results confirm that D614G and K856N mutations are key to explaining changes in spike flexibility within this region and, in context with simulation results, the increased number of RBDs in the up state in later VOCs relative to the Ancestral strain.

## Conclusions

We have used WE simulations to map the RBD opening pathways for Ancestral (novel 2019 strain), Delta, and Omicron BA.1 spike glycoproteins. We have observed RBD opening in the Ancestral strain spike is predicated upon breaking of the D614-K854 salt bridge interaction, but this salt bridge is reformed in up, open, and super-open RBD conformations. The D614G mutation abolishes the D614-K854 salt bridge, therefore removing the reliance of RBD opening on salt bridge breaking. We have also revealed an allosteric communication network between the NTD and RBD, also supported by Manrique et al,^74^ which passes through two linkers proximal to the 614 position. We see that mutation to G at the 614 position results in establishment of a new “lane” of communication within this allosteric network, thus indicating the D614 to K854 salt bridge likely disrupts optimal allostery within the S1 during RBD opening. Further, we hypothesize Omicron BA.1 spike’s N to K mutation at position 856 within the FPPR re-establishes a salt bridge to D568 nearby to the D614 to K854 salt bridge. We believe these D614G and N856K mutations represent goldilocks optimization of flexibility and stability around the allosteric communication pathways. We have also observed that the Omicron BA.1 RBD accesses a new “peel” conformation not yet seen in Ancestral or Delta spikes. This highly flexible peel state potentially indicates the “readiness” of the Omicron spike’s S1 domain to peel away from the S2 core, potentially leading to faster membrane fusion.

Taken together, these results support the presence of two linkers, the N2R and R2N, which facilitate allosteric communication within the spike’s S1. We believe the D614 to K854 salt bridge seen in Ancestral strain spikes causes steric congestion within the 614 proximal region preventing optimal communication via the R2N linker and instead only allowing communication via the N2R linker. The D614G mutation alleviates frustration in this area, allowing for concerted communication via both the N2R and R2N. We predict Delta and Omicron BA.1 spikes, both incorporating the D614G mutation, are likely to have much faster RBD opening rates, thus potentially contributing to increased propensities for these spikes to populate the 1-up, 2-up, and 3-up RBD states and increased ACE2 binding affinities. These findings are significant as loss of the D614-K854 salt bridge has long been scrutinized as the source of shifted RBD conformational landscapes in D614G and later variants, but the exact mechanism has yet to be so clearly described. Tracing the thermodynamic and kinetic fitness advantages of spike mutations aids in understanding viral fitness in humans and helps predict future gain of function mutations to this and other viral glycoproteins. Furthermore, observing structural changes between variant spike glycoproteins will enable the prediction of accessible or inaccessible antibody epitopes on variant spike surfaces. All in all, our work reveals the nature of the D614G mutation and its impact on spike structure, dynamics, and function relationships.

## Methods

### Weighted Ensemble Simulations

For a detailed description of spike model construction, including setup, equilibration, and the selection of described regions, please refer to the Supporting Information Methods Sections 1.0-1.1. Each spike structure was pre-equilibrated for a total of ∼72 ns, and 50 frames from these pre-equilibration simulations were used to launch the initial WE replicas for RBD opening simulation, starting from the closed/all-RBD-down spike conformation. All WE simulations were performed with the WESTPA^80^ software and GPU-accelerated Amber dynamics engine.^81–84^, using the CHARMM36m all-atom force field^85–87^ and CHARMM-modified TIP3P water model. Each protein was solvated in a simulation box of roughly 220 Å x 220 Å x 220 Å water molecules and150 mM NaCl. For all WE simulations, a fixed time interval of 100 ps was used with a target number of 8 trajectories/bin. Ancestral and Delta simulations were performed with Oracle Cloud Computing resources and Omicron simulations were performed on Delta at the National Computing for Supercomputing Applications (University of Illinois Urbana-Champaign).

### Pathway Clustering

To determine distinct pathway classes, a.k.a. routes, from each simulation, we applied the Linguistics Pathway Analysis of Trajectories with Histograms (LPATH) method^88^ as described in Supporting Information Methods Section 1.2.2.

### Dynamical Network Analysis Methods

To investigate the changes in conformational dynamics in the Spike protein across different variants, we employed a dynamical network approach using the routes extracted from complete weighted ensemble (WE) trajectories. The network was constructed by treating each residue’s Cα atom and each glycan’s C1 atom as a node. Edges between nodes were created based on a distance cutoff of 6 Å between the respective Cα/C1 atoms, and correlation values were derived from the WE pathway simulation data and were used to define the weight of each node and edge. The Weighted Implementation of Suboptimal Paths (WISP) method^77^ was applied to calculate the shortest paths between all residue pairs, and the Python library NetworkX,^89^ utilizing Dijkstra’s algorithm, was used for shortest path computation. Visualization was performed using VMD, showing the top 1000 most relevant edges and nodes. These dynamical networks were analyzed to examine the communication pathways across the protein and evaluate how they differ between variants. For complete WISP methods please Supporting Information Methods Section 1.2.4.

### Simulation Analysis Methods

All contact analyses, φ/ψ distributions, domain motions (distances, angles, dihedrals), root mean square fluctuations, were calculated via in-house scripts written with MDAnalysis.^90,91^ All analyses scripts are shared along with shared trajectories and resultant data files, as detailed in the Data and Code availability sections. For complete analysis details please Supporting Information Methods Sections 1.2.2, 1.2.5-1.2.8.

### HDXMS Methods

<u>VLP Prep:</u> VLP preparation methods were followed closely from previous work by Plescia et. al.,^92^ thus for complete details see that work. HEK293 cells were cultured in DMEM with 10% FBS and 1% penicillin-streptomycin at 37°C and 5% CO₂ until reaching 70-80% confluency. Cells were transfected with pcDNA3-M, pcDNA3-N, pCMV-E, and an S protein plasmid using PEI as a transfection reagent. After incubation, the media was replaced, and 48 hours post-transfection, the supernatant was collected and clarified through sequential centrifugation. The clarified media was then ultracentrifuged over a sucrose cushion to isolate VLPs, which were resuspended in TNE buffer and stored at −20°C. Samples were shipped on ice, flash frozen, and stored at −80°C before HDXMS analysis. <u>HDXMS labeling and LCMS:</u> VLP samples were incubated at 37°C for 3 hours before labeling. Incubated samples were diluted in labelling buffer prepared by diluting 20X TNE in D₂O (63% final D₂O concentration). Deuterium labeling was performed for 0, 1, 10, and 100 minutes, followed by quenching with a solution containing 4 M GdnHCl, 0.4 M TCEP, and 2.4 mM DDM. The labeled samples were digested online using a pepsin column, and peptides were separated via Waters UPLC with a controlled temperature and acetonitrile gradient. Mass spectra were acquired using a Waters SELECT SERIES Cyclic IMS-q-tof. <u>HDXMS analysis:</u> Peptides of WT and SARS-CoV-2 variant S were identified using PROTEIN LYNX GLOBAL SERVER (PLGS) with HDMSE mode for non-specific protease cleavage. Deuterium exchange was quantified using DynamX v3.0 with specific filtering criteria (minimum intensity=2000, minimum peptide length = 4, maximum peptide length =25, minimum products per amino acid + 0.2, and a precursor error tolerance of 10 ppm), and only peptides matching those criteria in 2 of 3 undeuterated replicates were included. Peptides unique to each S variant were identified and analyzed similarly for deuterium exchange. The data were processed in DynamX and visualized using MATLAB, with relative deuterium exchange reported as RFU, and the results will be deposited in the ProteomeXchange Consortium via PRIDE.

## Abbreviations

S1: spike domain 1 (residues 13 to 685)
S2: spike domain 2 (residues 686 to 1140)
NTD: N-terminal domain (residues 13 to 291)
RBD: receptor binding domain (residues 331 to 528)
RBM: receptor binding motif (residues 437 to 508)
SD1: spike subdomain 1 (residues 529 to 589)
SD2: spike subdomain 2 (residues 590 to 675)
FCS: furin cleavage site (residues 675 to 690)
FP: fusion peptide (residues 817 to 834)
FPPR: fusion peptide proximal region (residues 835 to 855)
HR1: heptad repeat 1 (residues 910 to 984)
CH: central helix (residues 985 to 1034)
CD: connecting domain (residues 1035 to 1068)
HR2: heptad repeat 2 (residues 1163 to 1210)
TM: transmembrane domain (residues 1214 to 1234)
CT: cytoplasmic tail (residues 1235 to 1273)
N2R: N-terminal Domain to the Receptor Binding Domain, linker
R2N: Receptor Binding Domain to the N-terminal Domain, linker
630loop: loop within the SD2 spanning residues 620-640
MD: molecular dynamics
WE: weighted ensemble
MAB: Minimal Adaptive Binning
LPATH: Linguistics Pathway Analysis of Trajectories with Histograms) method
WISP: Weighted Implementation of Suboptimal Paths (WISP) method
HDXMS: Hydrogen/Deuterium Exchange Mass Spectrometry

## Data availability

Selected WE simulation trajectories, along with input scripts and MDAnalysis scripts will be made available for download on the Amaro Lab website (https://amarolab.ucsd.edu/data.php#covid19) upon publication. An abridged set of input scripts has been uploaded for consideration by reviewers.

## Code availability

This work used standard builds of the WESTPA 2020.02 software (https://github.com/westpa/westpa) and Amber 18 (https://ambermd.org). Simulations were performed according to best practices for running WE simulations.^93^ MAB binning was used according to Torrillo et al.^70^

## Author Contributions

FLK, ATB, MKEB, MAR, LC, SHA, designed and conducted all weighted ensemble simulations. FLK, ATB, MKEB, CCT, LC, SHA, performed all computational analyses. AJG, SB, GA performed all HDXMS experiments and analysis. LTC, REA, SHA, GA guided and oversaw all simulations, experiments, and analyses.

## Acknowledgements

FLK, LC, LTC, SHA, and REA thank Prof. Jason McLellan for continued advice and conversations regarding SARS-CoV-2 spike conformational landscapes function. All authors would like to thank the gracious donation of compute time from Oracle Cloud Computing for Research and TACC Frontera for their never-ending support of our simulations. Compute resources for Omicron BA.1 simulations were supported by XSEDE (NSF TG-CHE060073) and ACCESS (NSF TG-CHE060063) allocations. We thank Robert Stahelin and Elijah Bass (Purdue University, IN) for the production of SARS-CoV-2 VLPs. We thank Thomas Wales and Bindu Srinivasu (Northeastern University, MA) for VLP HDXMS data collection.

