## Supporting Information for "D614G reshapes allosteric networks and opening mechanisms of SARS-CoV-2 spikes"

**Classification**: Biophysics and Computational Biology

**Keywords:** COVID19, SARS-CoV-2 spike glycoprotein, weighted ensemble simulations, receptor binding domain opening, ACE2 binding

### Table of Contents:

1. Methods:
    1.0 Computational Model Construction of Ancestral, Delta, and Omicron Spike Proteins
        1.0.1 Ancestral
        1.0.2 Delta
        1.0.3 Omicron BA.1
        1.0.4 Glycosylation/ Protonation/ Solvation/ Neutralization
        1.0.5 Equilibration via conventional Molecular Dynamics Simulations
        1.0.6 Equilibration via cMD in Amber
    1.1 WE simulation methods for Ancestral, Delta, and Omicron Spike Proteins
1.1.0 Installation of WESTPA and Amber on Oracle Cloud
    1.1.1 Installation of WESTPA and Amber on Frontera
        1.1.2 WE simulations
    1.2 Analysis methods
        1.2.0 Conformational ensembles of stable and transition states

1.2.1 Path ensembles for the spike-opening process
1.2.2 Clustering of successful pathways
1.2.3 Dynamical Network Analysis via WISP
1.2.4 Residue to Residue Contact Distance Measurements
1.2.5 Φ/Ψ distributions
1.2.6 Distance between 630 loop and spike core
1.2.7 Distances and areas between SD2, FPPR, and residues 590-594 within the R2N
2. Results and Discussion
2.0 Tables
**Table S1.** Total trajectory file sizes from full ensembles.
**Table S2.** Table outlining all HDXMS experimental conditions.
2.1 Figures
**Figure S1:** Distributions of relative molecular opening times for all spikes
**Figure S2:** The LPATH method identifies two major pathway classes present in simulations of all three spike variants.
**Figure S3:** Allosteric networks calculated from Routes 1 and 2 for all spikes.
**Figure S4:** Relative weight differences across allosteric networks.
**Figure S5:** Distance between each 630loop's center of mass and the center of mass of the spike core.
**Figure S6:** Minimum distance between $X614_A$ to $K854_B$.
**Figure S7:** Φ/Ψ distributions for residues 590 to 594.
**Figure S8:** Distance between N317-NH2 to F592-C=O.
**Figure S9:** Minimum distance between X856 to D568.
**Figure S10:** Distance between the SD2 and R2N centers of mass.
**Figure S11:** Distance between the R2N and FPPR centers of mass.
**Figure S12:** Distance between the FPPR and SD2 centers of mass.
**Figure S13:** Area between the SD2, R2N, and FPPR centers of mass.
**Figure S14:** Trends correlating with spike opening described in the main text shown for all successful stitched trajectories (SSTs) for Ancestral, Delta, and Omicron spikes.
**Figure S15:** Color scheme details.
3. References

### 1. Methods:

***1.0 Computational Model Construction of Ancestral, Delta, and Omicron Spike Proteins:*** Fully glycosylated, all-atom models of Ancestral, Delta, and Omicron BA.1 SARS-CoV-2 spike glycoprotein head domains (residues 13 to 1140) were constructed according to protocols described in the supporting information of Kim and Kearns et al, 2023.[1] A summary of model construction protocols will be described here.

<u>1.0.1 Ancestral:</u> Ancestral "closed"/all RBD down system was based on PDB ID 6VXX,[2] with fully resolved NTD, RBD, and pre-fusion loops grafted from PDB ID 7JJI.[3]

<u>1.0.2 Delta:</u> Delta "closed"/all RBD down system was constructed using the Ancestral "closed"/all RBD down structure as a baseline. Single point mutations were incorporated via the "mutate" command in VMD's psfgen.[4] Delta's significantly remodeled NTD was incorporated by grafting the NTD from a partially resolved Delta spike structure, PDB ID 7SO9.[5]

1.0.3 Omicron BA.1: Omicron "closed"/all RBD down system was based on PDB ID 7TF8.[6] Missing loops from the furin cleavage site were grafted in from PDB ID 6VSB.[7] The 7FT8 structure contains many missing loops, particularly in the NTD, thus we incorporated a fully-resolved Omicron BA.1 NTD structure from PDB ID 7K4N.[8]

1.0.4 Glycosylation/ Protonation/ Solvation/ Neutralization: All spike models were glycosylated following the same glycoprofile as used by Casalino et al.,[9] consistent with Watanabe et al.[10] Schrödinger Protein Preparation Wizard[11] does not handle N- or O-linked glycans, whereas standalone PROPKA3[12] is capable of treating them, thus we predicted the pKa baseline for each titratable residue with standalone PROPKA3 and predicted Histidine tautomeric states (i.e., HSD vs HSE) via Schrödinger's Protein Preparation Wizard. Spike models were then each solvated in explicit TIP3P[13] water boxes of 220 Å x 220 Å x 220 Å and neutralized with 150 mM NaCl.

1.0.5 Equilibration via conventional Molecular Dynamics Simulations: One replica per spike (3 closed spike models in total) was then minimized, heated, and equilibrated for 50 ns via conventional molecular dynamics (cMD) simulations in NAMD2.14 including under constant volume (NVT) and constant pressure (NpT) conditions,[14] according to the CHARMM36m all-atom force field,[15–17] and using TACC Frontera CPU resources. Standard MD simulation options were used including: timestep = 2 fs, temperature = 310 K, pressure = 1 atm, temperature and pressure maintained via the Nosé-Hoover thermostat[18] and Langevin barostat,[19] bonds to hydrogen constrained according to parameter values via the SHAKE algorithm,[20] long-range electrostatics handled via Particle Mesh Ewald[21] (interpolation order = 4 and grid spacing = 2 Å) with a 10-12-13.5 Å cutoff scheme. See Supporting Information Methods Section 2.1.1 in Kim and Kearns et al, 2023,[1] for complete details.

After this step, methods relevant to the current manuscript are no longer described in Kim and Kearns et al, 2023,[1] thus the following methods are unique to this work and thus described in full below.

1.0.6 Equilibration via cMD in Amber: Following the initial equilibration (50 ns) with NAMD2.14, CHAMBER[22] was used to convert CHARMM protein structure and CHARMM coordinate files to Amber topology and coordinate files, respectively, for each the spike protein systems. The Amber 22 software was then used to further energy minimize and equilibrate the system to prepare for WE simulations using WESTPA 2 (version 2022.03) and Amber 22. Energy minimization was performed in two stages. First the solvent and ions were minimized for 100,000 steps with harmonic position restraints on the solute heavy atoms. Next, the entire system was minimized without restraints. The system was equilibrated for 1 ns in the NPT ensemble with position restraints on the solute heavy atoms. Finally, an unrestrained production simulation was run for 20 ns. Temperature and pressure were maintained via a weak Langevin thermostat[18] (collision frequency of 1 $ps^{-1}$) and Monte Carlo barostat[19] (pressure changes attempted every 0.2 ps) at 300 K and 1 atm, respectively. Bonds to hydrogen were constrained to their equilibrium values via the SHAKE algorithm to enable a 2-fs time step.[20] Long-range electrostatics handled via Particle Mesh Ewald[21] with nonbonded, short-range interactions truncated at 10 Å. Conformations were saved every 0.01 ns. To generate an ensemble of equally-weighted closed-state conformations for initiating WE simulations of spike opening, we selected 50 conformations from the last 5 ns of the production simulation.

##### *1.1 WE simulation methods for Ancestral, Delta, and Omicron Spike Proteins:*

1.1.0 Installation of WESTPA and Amber on Oracle Cloud:

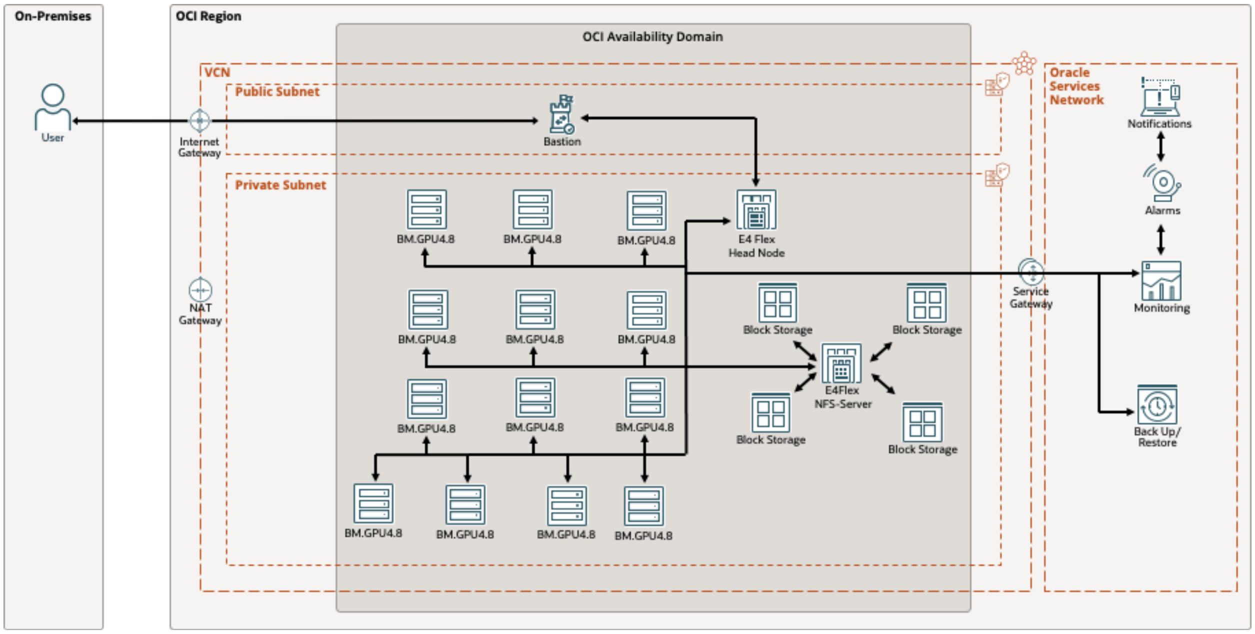


A custom OCI image was created using Oracle-Linux-7 as the base, and all required drivers, simulation package Amber 22 and AmberTools were installed following the installation instructions on the manual. WESTPA 2 was installed using Python pip. As described in the architecture diagram, using the custom image and a BM.4.8 instance configuration, a cluster of A100 GPUs was launched on the OCI tenancy private subnet using the OCI-instance pool. As a head node, an AMD E.4 Flex instance was launched. A custom network file system (NFS) was constructed by connecting multipath enabled ultra-high-performance block volumes into an E4.Flex instance running an NFS server where the number of service threads equals the number of serving GPU instances. Computing resource management was done via simple Linux utility for resource management (SLURM). Storage input-output (I/O), NFS-server, and GPU node performance were monitored using the OCI-monitoring service. Noncritical performance data were sent to the user via OCI-notification email, while critical information was sent via OCI-notification short-message service (SMS). Finally, the on-premises user was given cluster access via a bastion server installed on the public subnet. Upon completion of the simulation, the GPU cluster was terminated while leaving the storage intact to reduce costs.

1.1.1 Installation of WESTPA and Amber on NCSA Delta: WESTPA 2 (2022.03) and Amber 22 were installed on TACC Frontera and NCSA Delta according to standard installation protocols.[22–24]

1.1.2 WE simulations: The Ancestral weighted ensemble method (WE) simulations were run for 600 WE iterations over 24.6 days using 112 NVIDIA A100 GPUs on Oracle Cloud, yielding an aggregate of 60.3 µs simulation time. These simulations generated 128 pathways from the closed to the up state, 69 pathways from the closed to the open state, and 25 pathways from the closed to the super-open state. The Delta WE simulations were run for 270 WE iterations over 11 days using 112 NVIDIA A100 GPUs on the same computing resource, yielding an aggregate of 28.9 µs of simulation time. These simulations generated 177 pathways from the closed to the up state, 141 pathways from the closed to the open state, and 15 pathways from the closed to the super-open state. The Omicron WE simulations were run for 400 WE iterations, yielding 35.5 µs of aggregate simulation time. These simulations were run for over 23.3 days in NCSA Delta multiple A100 GPUs. These simulations generated 184 pathways from the closed to the up state, 192 pathways from the closed to the open state, and 79 pathways from the closed to the super-open state.

##### *1.2 Analysis methods:*

1.2.0 Conformational ensembles of stable and transition states: The closed state ensemble consisted of structures with RBD-core distance < 55 Å. The ensemble of transition states between closed and up states consisted of structures with RBD-core distances 55 – 68 Å. The up-state ensemble consisted of structures with RBD-core distances 68 – 72 Å. The ensemble of transition states between up and open states consisted of structures with RBD-core distances 72 – 78 Å. The open state ensemble consisted of structures with RBD-core distances 78 – 82 Å. The ensemble of transition states between open and super-open states consisted of structures with RBD-core distances 82 – 85 Å. The super-open state ensemble consisted of structures with RBD-core distances >85 Å.

1.2.1 Path ensembles for the spike-opening process: The successful pathways that reached the up state (RBD-core distances 68 – 72 Å), the open state (RBD-core distances 78 – 82 Å), or the super-open state (RBD-core distances >85 Å) were obtained by counting all arrivals to that particular state at every WE iteration. We consider these pathways to be statistically independent pathways, as we had previously justified in Sztain et al (2021).[25]

1.2.2 Clustering of successful pathways: To determine distinct pathway classes from each simulation, we utilized the LPATH (Linguistics Pathway Analysis of Trajectories with Histograms) method.[26] The LPATH method involves three steps: discretization, extraction, and matching. The successful pathways mentioned in section 1.2.1 were used as input, with pathways from each simulation analyzed separately.

To discretize pathways, we clustered conformations using their using their two-dimensional progress coordinate values. Due to the large number of conformations, hierarchical clustering on the entire dataset was computationally costly. Instead, we performed hierarchical clustering (average linkage, distance threshold of 5) on conformations from the last 50 iterations of WE (last frame of each trajectory segment). This yielded 26 clusters for the Ancestral spike, 28 for the Delta spike, and 23 for the Omicron BA.1 spike. Following hierarchical clustering, we calculated the centroid of each cluster and assigned each conformation to its nearest centroid. These final 50-iteration conformations and cluster IDs served as training data for a *k*-nearest neighbors (KNN) classifier (k = 5). The trained KNN model then assigned cluster labels to the remaining conformations, ensuring consistency with hierarchical clustering results.

After discretizing the conformational space, we extracted trajectory segments for each spike variant that transitioned from an RBD-core distance of 55 to 80, sampling every other frame.

Next, we matched trajectories using a Gestalt pattern matching algorithm with a length correction described in the original LPATH paper.[26] Consecutive visited states were not condensed (consecutive states removed), but consecutive *pairs* were removed. Similarity scores between pathways were used to perform hierarchical clustering with the default linkage in the LPATH package. The resulting dendrograms, representing pathway similarities, were then divided into two pathway classes (routes).

1.2.3 Dynamical Network Analysis via WISP: To investigate the conformational dynamics of the Spike protein across different variants, we employed a dynamical network analysis approach using data from Weighted Ensemble (WE) simulations. First, from the pathway clustering trajectories (see Section 1.2.2), distance and cross-correlation matrices were computed using the *cpptraj* module of Amber 22,[23] considering the Cα atoms of protein residues and the C1 atoms of glycans. The cross-correlation matrix, calculated using Pearson's correlation coefficient, captures residue pair correlations by measuring the linear relationship between atomic motions. The correlation coefficients values range from -1 to 1. A value of +1 indicates fully correlated motion (residues moving in the same direction), 0 represents uncorrelated motion, and -1 corresponds to fully anti-correlated motion (residues moving in opposite directions).

By using the Weighted Implementation of Suboptimal Paths (WISP) method,[27] a network representation of the Spike protein was then constructed by mapping the *N* residues and glycans into *N* nodes of a weighted graph. Each residue and glycan were represented as a node, centered on its Cα/C1 atom, and an edge was established between two residues if their mean distance remained below 6 Å throughout the simulation. The edges were weighted based on the normalized cross-correlation matrix values using the transformation *(dij=-log |Cij|)*, where *Cij* is the cross-correlation coefficient between residues *i* and *j*, and *dij* represents a functional correlation-based distance. This transformation ensures that strongly correlated residue pairs have shorter edge distances, reflecting the probability of information transfer based on their correlation.

To identify key communication pathways, Dijkstra's algorithm was applied by using the NetworkX Python library.[28] This algorithm determines the shortest paths between residues by prioritizing edges with shorter functional distances, highlighting the most frequently used and highly correlated pathways. The method identifies edges that contribute most to allosteric communication and residues that are central to network connectivity. The resulting dynamical networks were visualized using VMD,[4] displaying the top 1000 most relevant edges and nodes based on their contribution to communication pathways. Finally, comparative analyses were conducted across different Spike protein variants to assess alterations in network topology and inter-residue communication, allowing us to investigate potential structural and functional changes induced by mutations.

To assess differences in residue-specific weights between variants, we conducted a comparative analysis of the two important regions, N2R and R2N (see Figure S4). For each residue, we calculated the total contribution of all its connected edges by summing the weights associated with that residue within these domains and then compared the resulting summed weights across different variants. For visualization, comparative plots were generated by computing the difference in summed weights between variants and the Ancestral protein. The Delta-Ancestral and Omicron-Ancestral differences were determined by subtracting the Ancestral residue weights from the corresponding residue weights in the Delta and Omicron variants, respectively.

1.2.4 Residue to Residue Contact Distance Measurements: MDAnalysis[29,30] was used to calculate per-residue contacts for every frame in each simulation and across all chains. The minimum distance between heavy atoms (not hydrogen) was used to calculate the contact distances for X614 to K854 and for X856 to D568. Since we were specifically interested in a side chain to backbone interaction in the case of N317 to F592, here we calculated the distance between N317's-NH2 and F592's backbone carbonyl oxygen. Contact distances were calculated for all complete ensemble trajectories, for stitched successful pathways, and for pathway classes (routes 1 and 2).

1.2.5 Φ/Ψ distributions: Ramachandran plots (Φ/Ψ distributions) were calculated per frame using MDAnalysis's Ramachandran class in the "dihedrals" module, for amino acids 589 to 594 from full ensemble trajectories, stitched successful pathways, and pathway classes (routes 1 and 2) for Ancestral, Delta, and Omicron spike proteins.

1.2.6 Distance between 630 loop and spike core: Distances between the 630loop and spike core were calculated per frame with MDAnalysis by selecting the center of mass of the 630loop's $C_\alpha$ atoms (residues 620 to 640) and the center of mass of spike central helices' $C_\alpha$ atoms (residues 747-784, 946-967, and 986-1034). Distances were calculated from full ensemble trajectories, stitched successful pathways, and pathway classes (routes 1 and 2) for Ancestral, Delta, and Omicron spike proteins.

1.2.7 Distances and areas between SD2, FPPR, and residues 590-594 within the R2N: Distances between the SD2, FPPR, and residues 590-594 within the R2N were calculated per frame with MDAnalysis by selecting the center of mass of the SD2's $C_\alpha$ atoms (residues 590-675, 691-697), the center of mass of the neighboring chain's FPPR's $C_\alpha$ atoms (residues 835-855), and the center of mass of a portion of the R2N's $C_\alpha$ atoms (residues 590-594). Distances and areas were calculated from full ensemble trajectories, stitched successful pathways, and pathway classes (routes 1 and 2) for Ancestral, Delta, and Omicron spike proteins.

##### *1.3 SARS CoV-2 experimental HDXMS methods:*

1.3.0: Expression and purification of SARS-CoV-2 VLPs (Ancestral, D614G, and Omicron): Human embryonic kidney (HEK) 293 cells (American Type Cell Collection, Manassas, VA) were maintained at 37°C and 5% $CO_2$ in Dulbecco's Modified Eagle's Medium (Thermo Fisher Scientific) supplemented with 10% fetal bovine serum (Biowest USA) and 1% penicillin streptomycin (Thermo Fisher Scientific). HEK293 cells were grown to 70-80% confluency in 100mm round dishes. pcDNA3-M, pcDNA3-N, pCMV-E, and an S protein plasmid (pCMV-wt S, pCAGGS-D614G S, or pCAGGS-BA.1 S) were transfected at a ratio of 1:1:1:2 using a 3:1 ratio of polyethyleneimine hydrochloride (PEI) (Polysciences) to DNA. The DNA and PEI were mixed in Opti-MEM (Fisher Scientific) and incubated at room temperature for 10-15 minutes before being added dropwise to cells. The cells were incubated at 37°C and 5% $CO_2$. The cell media was replaced with fresh DMEM supplemented with 10% fetal bovine serum and 1% penicillin streptomycin after five to six hours post-transfection. At 48 hours post-transfection, the media was collected in a falcon tube containing 0.1x HaltTM Protease Inhibitor Cocktail (Thermo Fisher Scientific). The media was clarified by centrifugation at 1000 x *g* for 10 minutes at 4°C, transferred to a new falcon tube, and further clarified by centrifugation at 2000 x *g* for 10 min at 4°C. The supernatant was then loaded onto a 20% sucrose cushion in TNE buffer (50 mM Tris-HCl, 100 mM NaCl, 0.5 mM EDTA, pH = 7.4) and ultracentrifuged at 100,000 x *g* in a Beckman Type 70 Ti rotor for 3 hours at 4°C. VLP pellets were dried, gently resuspended in TNE buffer, and stored at -20°C before shipping. Samples were shipped on ice then flash frozen and stored at -80°C before HDXMS analysis.

1.3.1: Deuterium exchange of VLP on cyclic-IMS-Q-Tof: Labeling buffer was prepared by diluting 20 X TNE in $H_2O$ in $D_2O$ (99.9%). VLP and recombinant samples were incubated at 37 ℃ for 3 hours before labeling to ensure trimer splaying was not observed (Costello et al. *Nat. Struct. Mol. Biol*. 2022), (Edwards et al. *Nat. Struct. Mol. Biol.* 2021). 20 µL of sample were added to 40 µL of labeling buffer for a final labeling concentration of 63.3%. Deuterium labeling was conducted for 1, 10, and 100 min at 20°C for VLPs and 10 min at 20°C for recombinant S. After labeling, 20 µL of prechilled quench solution (4 M Guanidinium Hydrochloride, 0.4 M Tris(2-carboxyethyl) phosphine, 2.4 mM n-Dodecyl β-D-Maltoside was added to the 60 µL labeling mixture to bring the reaction to pH 2.5.

1.3.2: LC and Mass spectrometry of VLP on cyclic-IMS-Q-Tof: Deuterated and control samples were digested online at 15 °C using an AffiPro pepsin column (AffiPro, AP-PC-001). Peptides were trapped and desalted on a VanGuard Pre-Column trap [2.1 mm × 5 mm, ACQUITY UPLC BEH C18, 1.7 µm (Waters, 186002346)] for 3 minutes at a flow rate of 100 µL/min. Elution was achieved using a 5%–35% acetonitrile gradient over 10 minutes at 100 µL/min, with separation on an ACQUITY UPLC HSS T3 column [1.8 µm, 1.0 mm × 50 mm (Waters, 186003535)]. Mass spectra were acquired on a Waters SELECT SERIES Cyclic IMS instrument coupled with a UPLC I-Class system and HDX manager. Spectra were recorded over an m/z range of 50–2000 with the following settings: capillary voltage, 3.0 kV; trap collision energy, 4 V; sampling cone, 40 V; transfer CE ramp, 15–50 V; source temperature, 80 °C; and desolvation temperature, 450 °C. Ion mobility separation was performed with a single pass using a sequence of 10 ms injection, 3 ms separation, and 34 ms ejection/acquisition.

1.3.3: Peptide identification and hydrogen-deuterium exchange analysis: Peptides of ancestral and SARS CoV-2 variant S were identified through independent searches of mass spectra from the undeuterated samples. Peptides common to ancestral and variant S were identified from a database containing the amino acid sequence of D614G S using PROTEIN LYNX GLOBAL SERVER version 3.0 (Waters, Milford, MA) in HDMSE mode for non-specific protease cleavage. Search parameters in PLGS were set to "no fixed or variable modifier reagents" and variable N-linked glycosylation. Deuterium exchange was quantified using DynamX v3.0 (Waters, Milford, MA) with cutoff filters of minimum intensity = 2000, minimum peptide length = 4, maximum peptide length = 25, minimum products per amino acid = 0.2, and precursor ion error tolerance <10 ppm. Three undeuterated replicates were collected for WT and variant S, and the final peptide list includes only peptides that fulfilled the above-described criteria and were identified independently in at least 2 of the 3 undeuterated samples. Deuterium exchange in these peptides was analyzed using DynamX 3.0 with identical parameters described above. To analyze cyclic IMS data, PLGS and DynamX were modified as described in (Griffiths et. al. *Anal. Chem.* 2024). The average number of deuterons exchanged in each peptide was calculated by subtracting the centroid mass of the undeuterated reference spectra from each deuterated spectra. Peptides were independently analyzed for quality across technical replicates. The data for the uptake plots were acquired from DynamX v3.0 and plotted in MATLAB 2024a, The MathWorks Inc, Natick, MA, USA. Relative deuterium exchange plots are reported as RFU which is the ratio of exchanged deuterons to possible exchange deuterons. The mass spectrometry proteomics data will be deposited to the ProteomeXchange Consortium via the PRIDE partner repository.

### 2. Results and Discussion:

##### *2.0 Tables:*

**Table S1. Total trajectory file sizes from full ensembles, all values in GB.** Note: all water and ion atoms have been stripped prior to estimating file size, complete data sets with solvent atoms are larger.

| | Ancestral | Delta | Omicron BA.1 |
|---|---|---|---|
| **Closed** | 164 | 36 | 25 |
| **Closed to Up** | 431 | 216 | 137 |
| **Up** | 22 | 22 | 26 |
| **Up to Open** | 26 | 47 | 75 |
| **Open** | 4.7 | 13 | 19 |
| **Open to Superopen** | 8.2 | 17 | 35 |
| **Superopen** | 1.5 | 0.69 | -- |
| **Total** | 657.4 | 351.7 | 317 |

**Table S2.** Table outlining all HDXMS experimental conditions.

| **Spike Dataset** | VLP ancestral on cyclic | VLP D614G on cyclic | VLP Omicron BA.1 on cyclic |
|---|---|---|---|
| **HDX reaction details** | Labeling buffer: Tris-NaCl-EDTA buffer (TNE) pH 7.4 prepared in 99.9% $D_2O$ to a final concentration of 94.9% $D_2O$<br>Quench buffer: 4 M Guandinium hydrochloride, 0.4 M TCEP, 1.4mM DDM<br>HDXMS reaction: 40 µL Labeling buffer added to 20 µL VLP S sample for deuterium exchange at 20°C and quenched with 20 µL of quench buffer by bringing reaction to pH 2.5 at 0°C. | | |
| **Spike incubation** | 3h at 37°C | | |
| **HDXMS un-deuterated controls** | 40 µL of TNE pH 7.4 was added to 20 µL of sample (either concentrated VLP solution or diluted recombinant sample) and 20 µL of quench buffer was added by bringing reaction to pH 2.5 at 0°C. | | |
| **HDXMS time course (min)** | 1, 10, 100 | | |
| **Number of peptides** | 55 | | |
| **Sequence coverage** | 45.6% | | |
| **Peptide redundancy** | 1.29 | | |
| **Replicates** | 3 technical replicates were acquired for all time points and states. | | |

##### *2.1 Figures:*

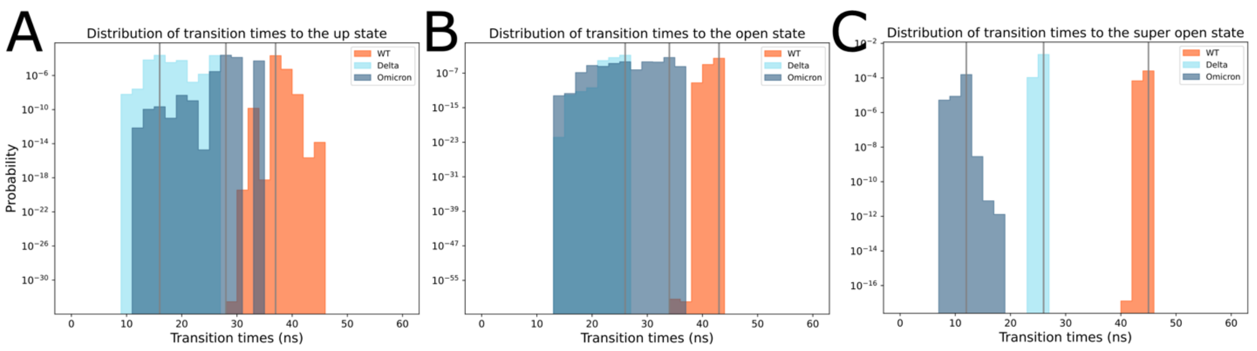

**Figure S1: Distributions of relative molecular opening times** observed for Ancestral (WT), Delta, and Omicron BA.1 spike proteins from the closed to (A) up, (B) open, and (C) superopen states. Ancestral/WT, Delta, and Omicron BA.1 distributions are given in orange, cyan, and blue histograms respectively.

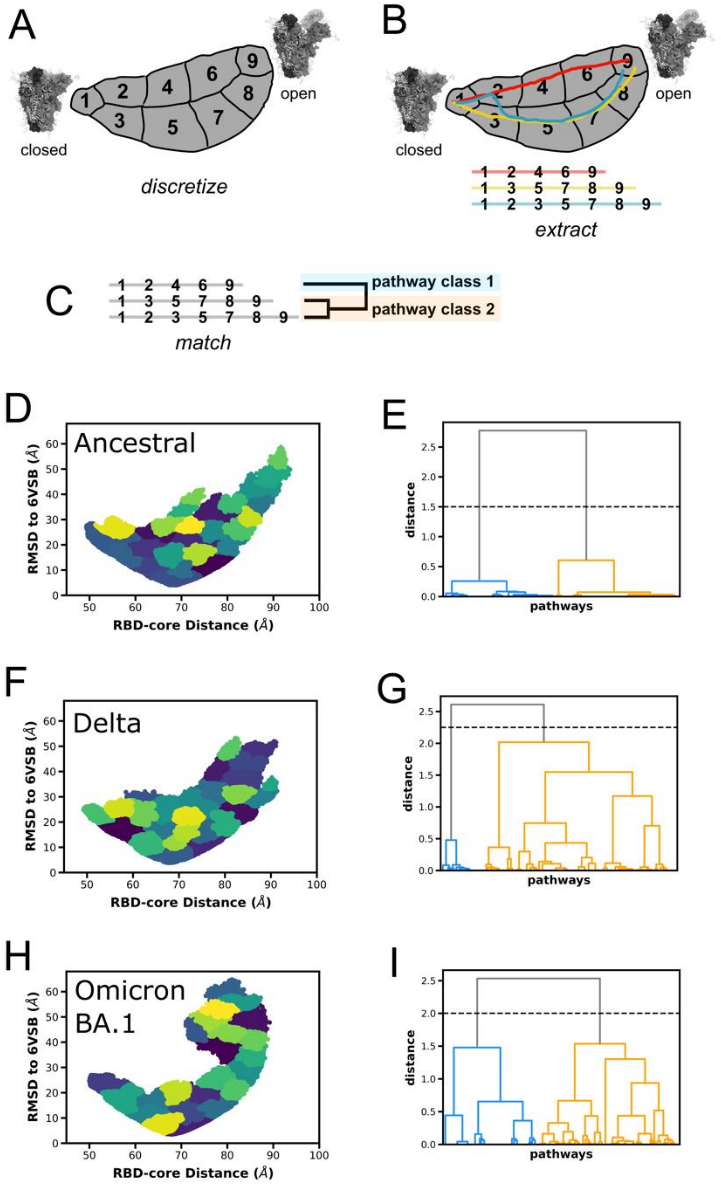


**Figure S2: The LPATH method identifies two major pathway classes present in simulations of all three spike variants.** (A) The sampled conformational space between two end state conformations (closed and open) is discretized into clusters and each conformation is assigned to a cluster. (B) Three pathways that have successfully left the closed state and reached the open state are extracted, with their cluster assignments serving as labels. (C) Strings of the states visited by each of the extracted pathways are matched with hierarchical clustering, revealing that, in this example, the yellow and blue pathways from panel B are more related to each other than to the red pathway. Panels (D), (F) and (H) show the discretized state space for the three spike variants as a function of the two-dimensional progress coordinate used for the WE simulations. The different colors represent different clusters determined through the clustering protocol (described above). Panels (E), (G) and (I) show, for each state space discretization, the resulting dendrogram that results from hierarchical clustering of the extracted pathways. Each dendrogram was divided into two pathway classes.

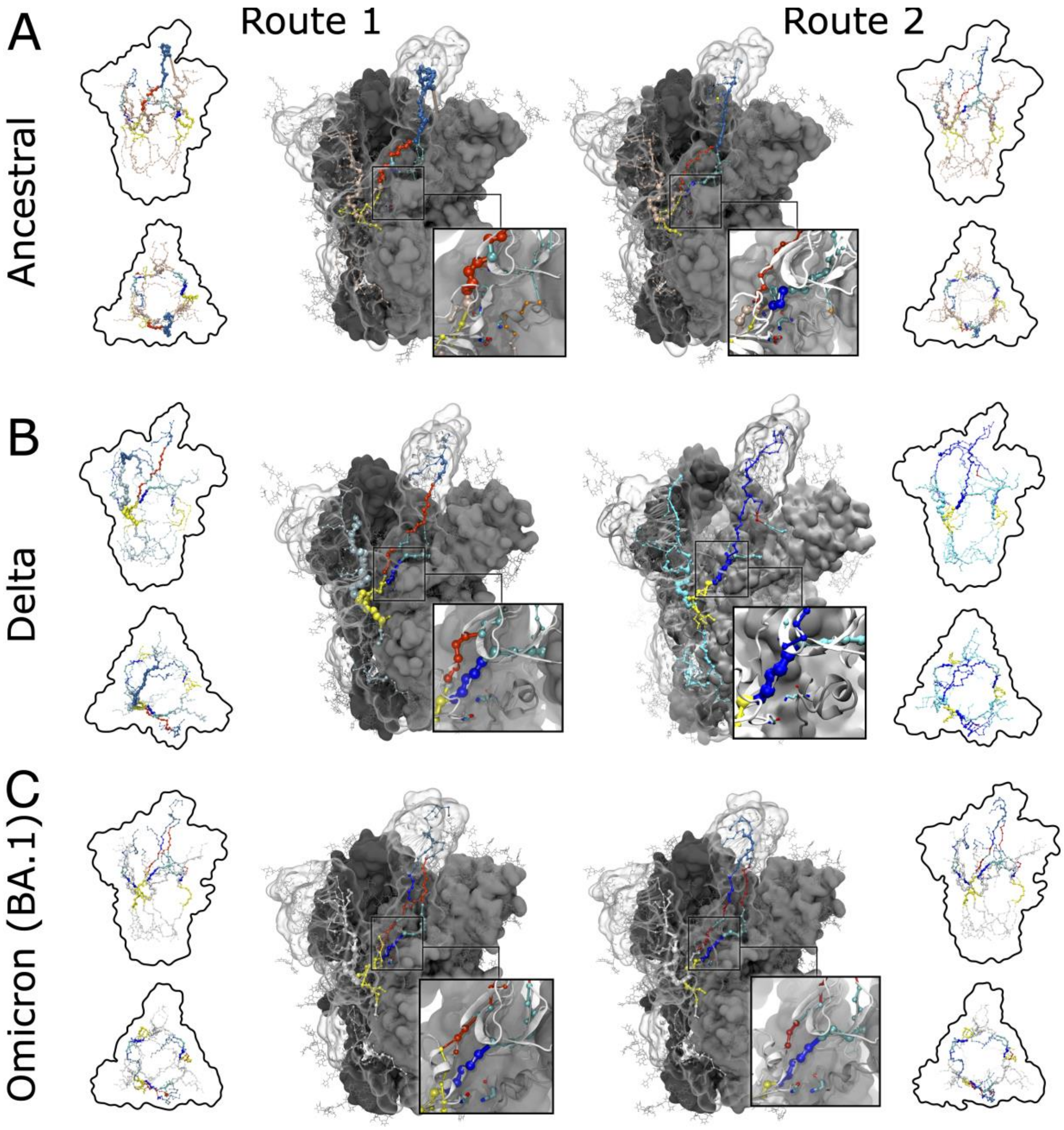


**Figure S3: Allosteric networks calculated from Routes 1 and 2** for (A) Ancestral, (B) Delta, and (C) Omicron BA.1 spikes. Panels also show side and top views of full networks within silhouette of spike structure to show correlated motions between spike monomers. Insets depict zoom ins of the 614 proximal region per route and per spike.

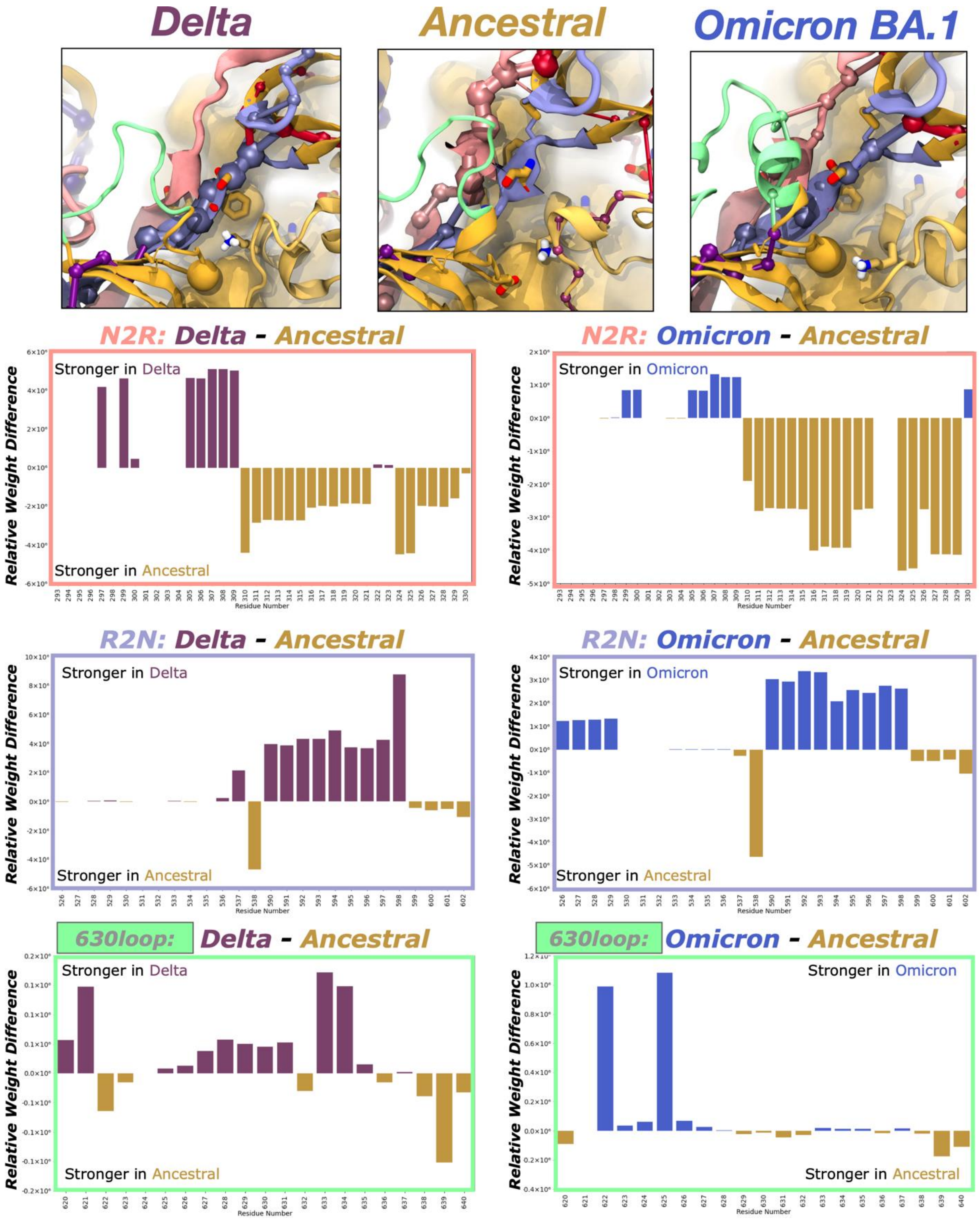


**Figure S4: Relative weight differences across allosteric networks** between Ancestral and Delta, and Ancestral and Omicron allosteric networks for amino acids within the N2R, R2N, and 630loop regions (calculated for Ancestral spike's route 1, Delta spike's route 2, and Omicron spike's route 1).

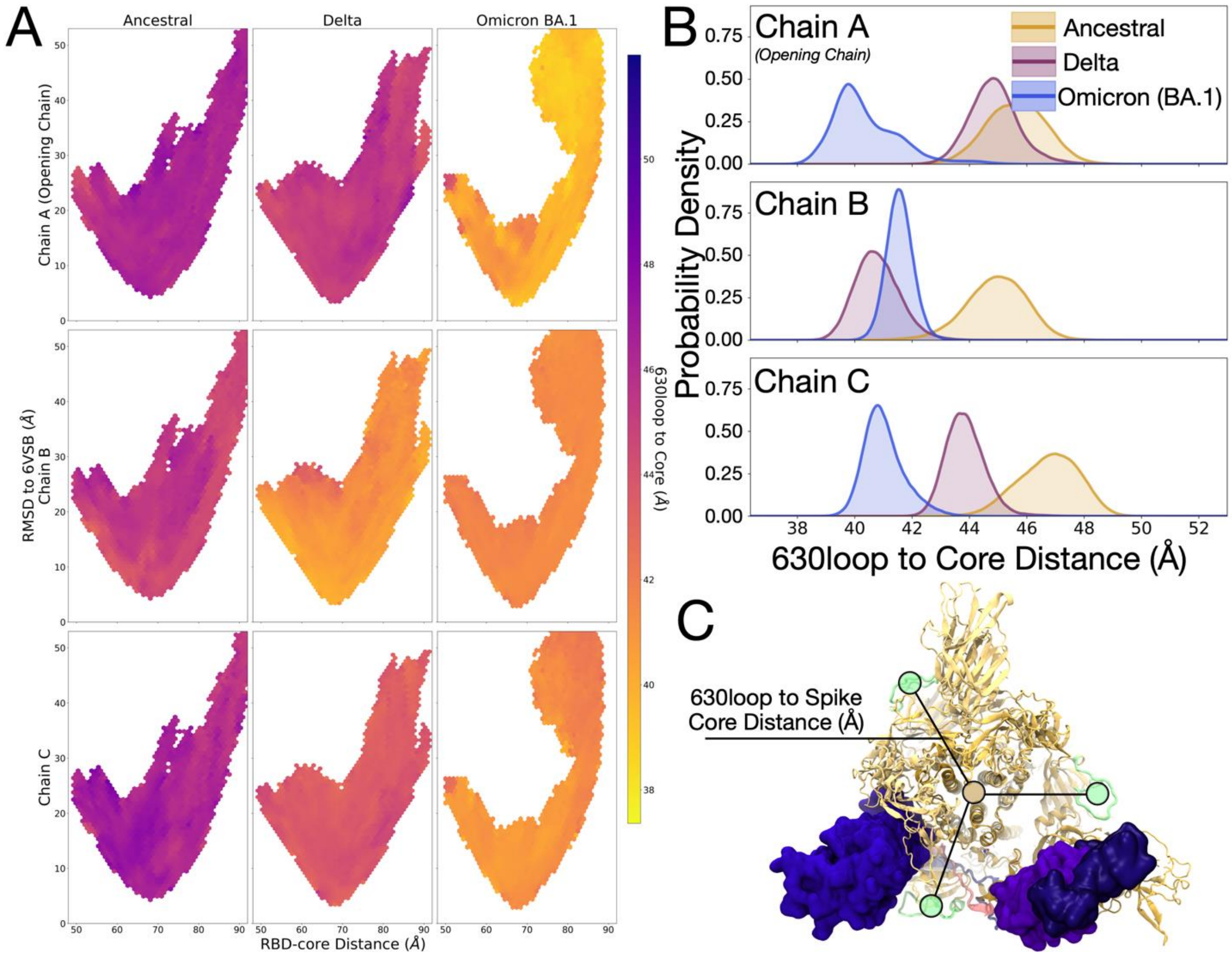


**Figure S5: Distance between each 630loop's center of mass and the center of mass of the spike core** (central helices). Data are shown as (A) histograms plotted as a function of the two-dimensional progress coordinates used to facilitate WE simulations and colored according to the average 630loop to Core Distance in Å and (B) one dimensional kernel probability density plots reporting the likelihood of 630loop to Core Distances as a function of spike protein and chain identity.

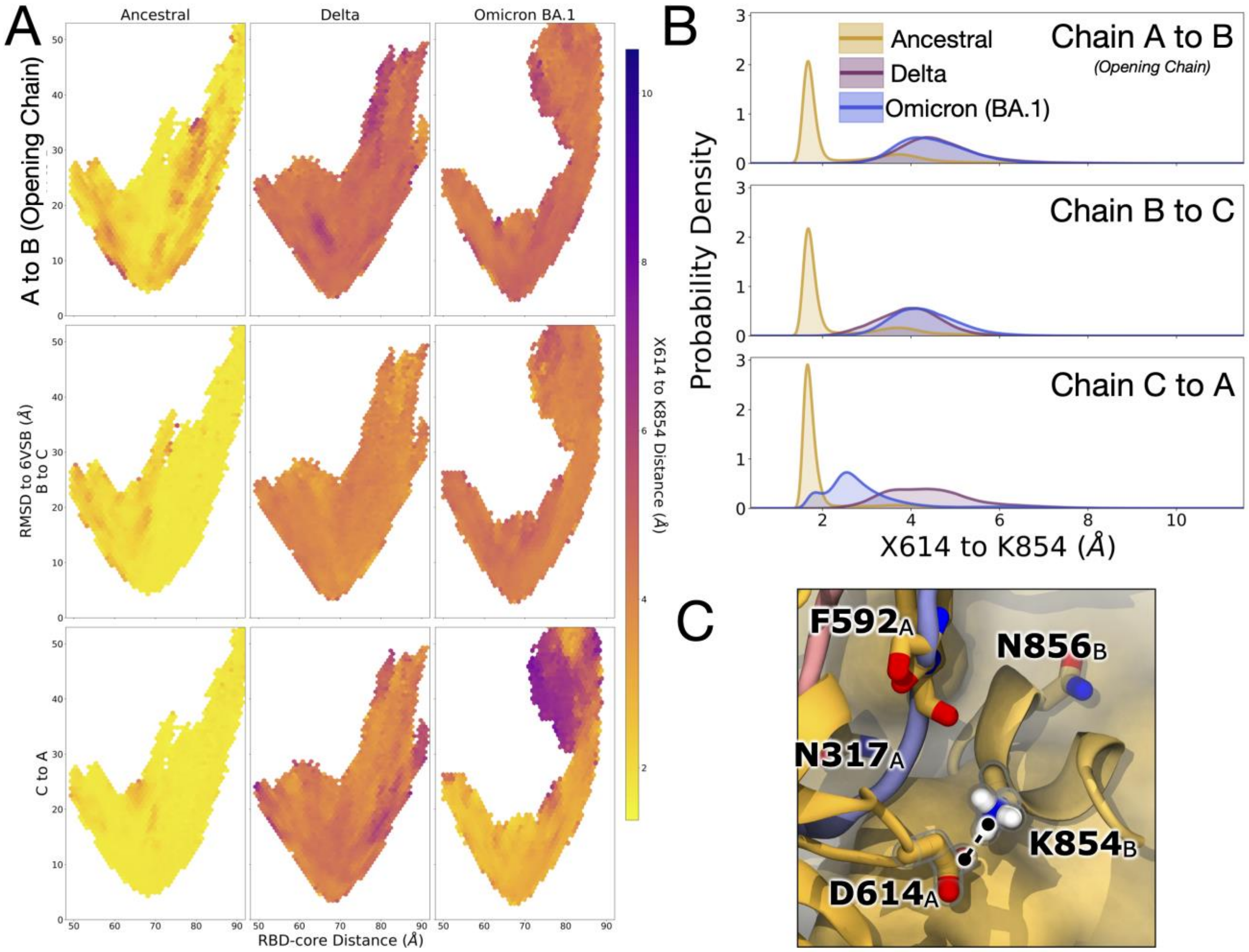


**Figure S6: Minimum distance between X614$_A$ to K854$_B$** for (A) complete ensemble sets, (B) stitched successful pathways, and (C) clustered pathway classes (routes) 1 and 2 per Ancestral, Delta, and Omicron BA.1 spikes during RBD opening. All data is plotted as 2D hexagonal histograms as a function of the 2D progress coordinate – i.e., RBD-Core Distance (Å) by RMSD to 6VSB (Å) -- and colored according to the average minimal distance between X614$_A$ to K854$_B$ per bin.

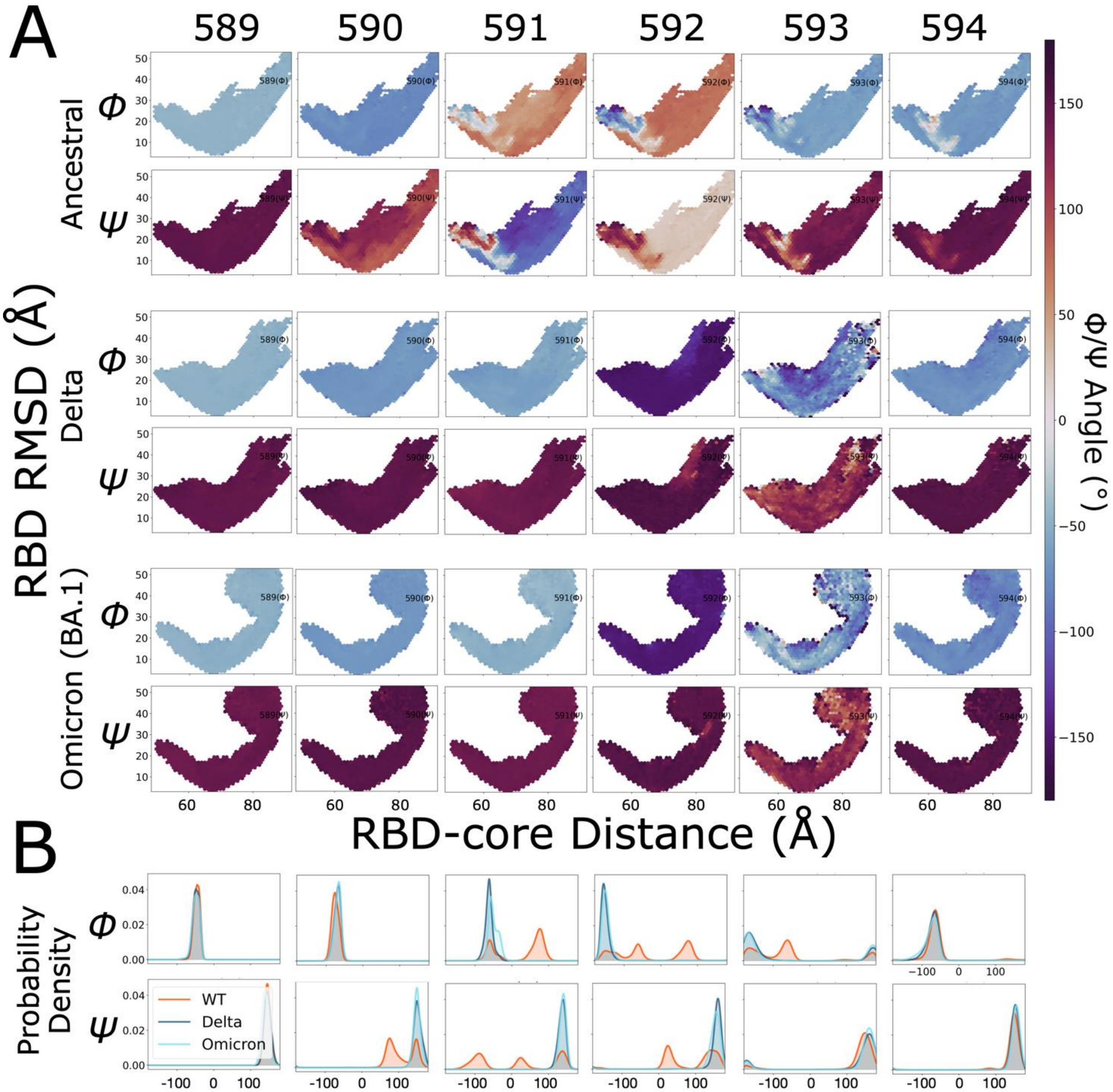


**Figure S7: Φ/Ψ distributions for residues 590 to 594** on the Chain A protomer, the RBD-opening chain, plotted as (A) histograms as a function of the 2D progress coordinates used to drive WE simulations, i.e., RBD-core Distances (Å) and RMSD to 6VSB (Å), colored according to average φ/ψ angle per residue per bin, and (B) one dimensional kernel probability density plots reporting the likelihood of φ/ψ distributions as a function of spike protein residue identity.

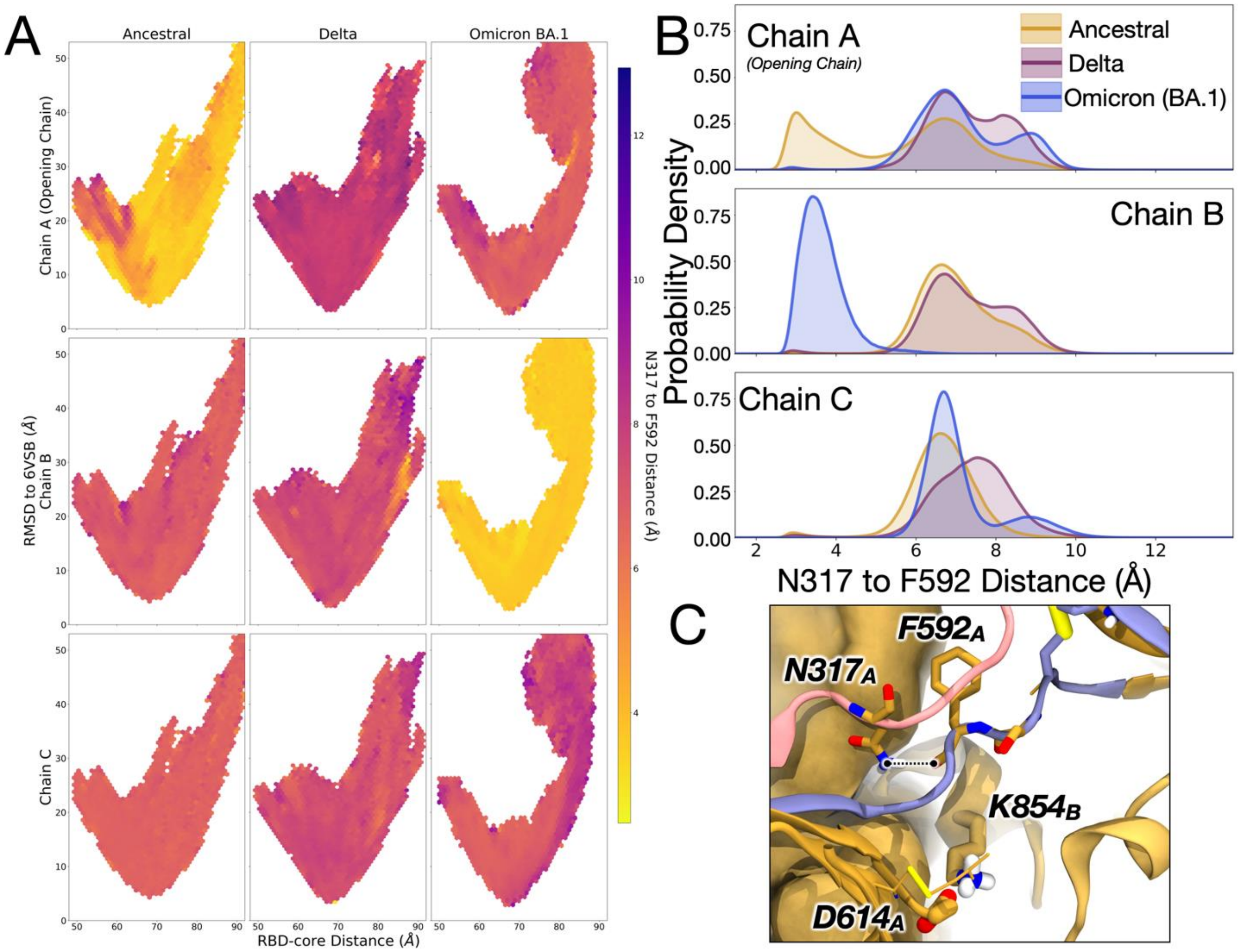


**Figure S8: Distance between N317-NH2 to F592-C=O** for chains A (opening), B, and C, shown as (A) hexagonal histograms as a function of the 2D progress coordinates used to run WE simulations, i.e., RBD-Core Distances (Å) and RMSD to 6VSB (Å), and colored according to the average distance for structures within each bin, and (B) one-dimensional distributions as a function of distance between N317's-NH2 to F592's-C=O. Panel (C) highlights the distance calculated herein between N317-NH2 to F592-C=O.

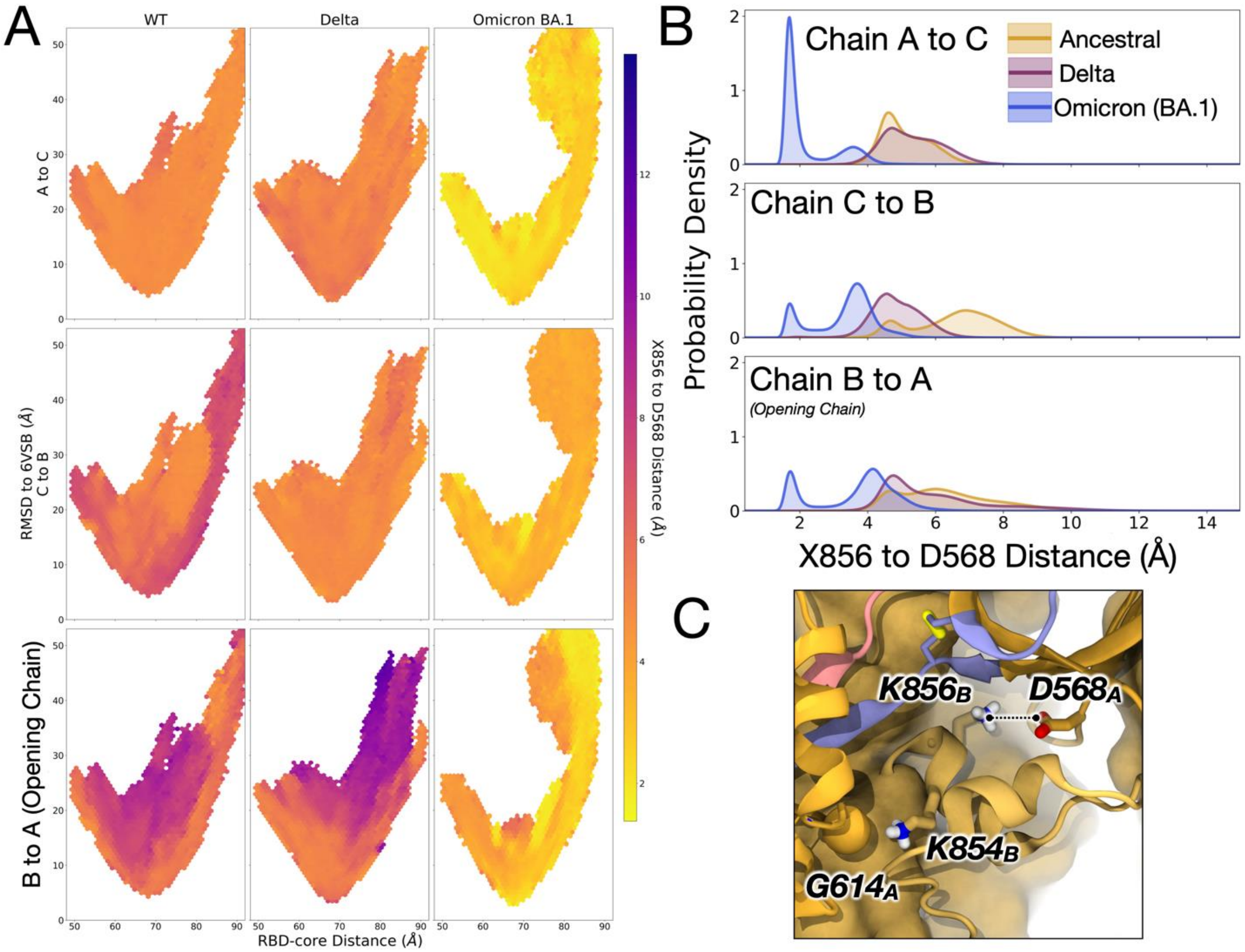


**Figure S9: Minimum distance between X856 to D568** for chains A-B (opening), B-C, and C-A, shown as (A) hexagonal histograms as a function of the 2D progress coordinates used to run WE simulations, i.e., RBD-Core Distances (Å) and RMSD to 6VSB (Å), and colored according to the average minimum distance for structures within each bin, and (B) one-dimensional distributions as a function of distance between X856 to D568. Panel (C) highlights the distance calculated herein between X856 to D568.

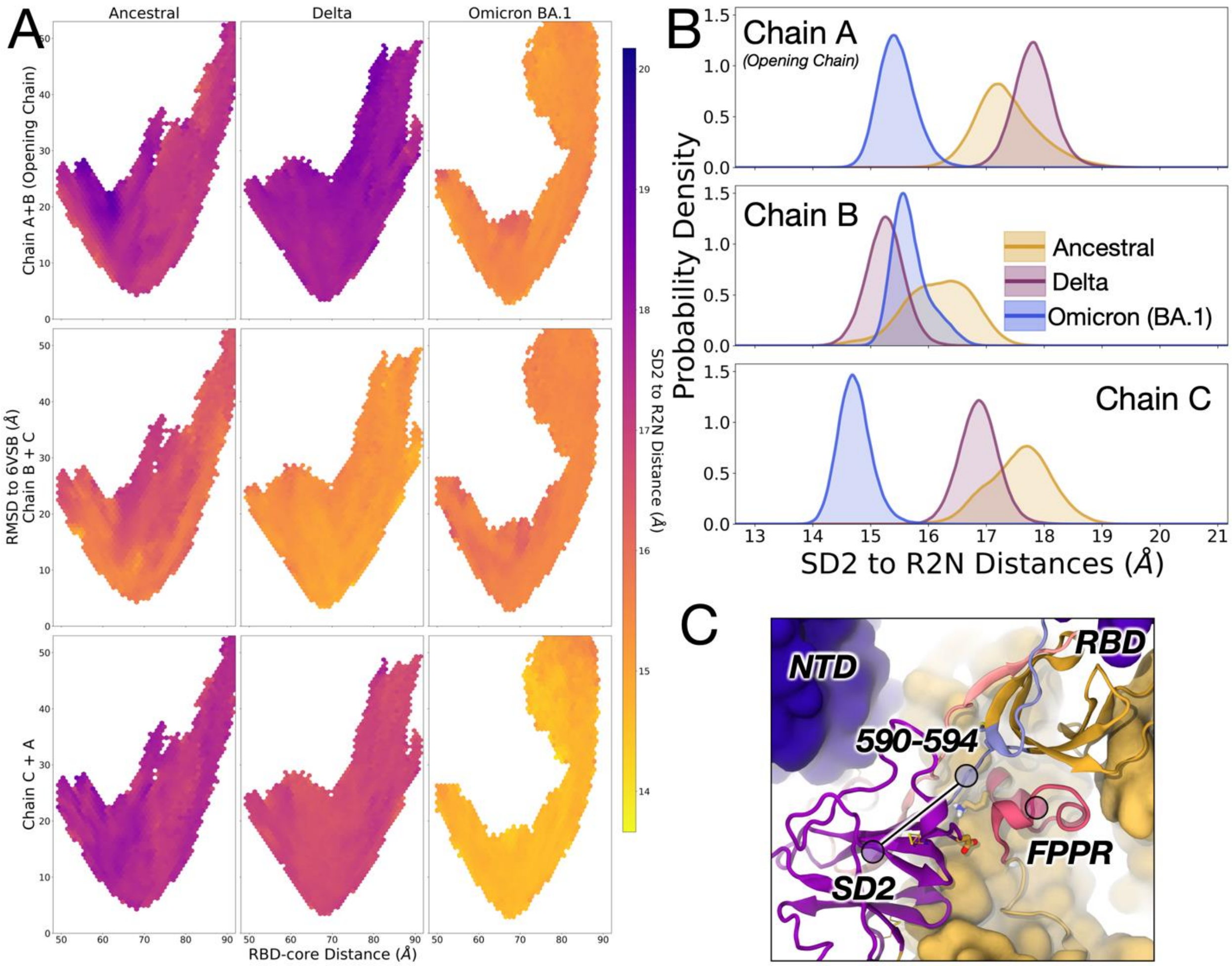


**Figure S10: Distance between the SD2 and R2N centers of mass** for chains A (opening), B, and C, shown as (A) hexagonal histograms as a function of the 2D progress coordinates used to run WE simulations, i.e., RBD-Core Distances (Å) and RMSD to 6VSB (Å), and colored according to the average distance for structures within each bin, and (B) one-dimensional distributions as a function of distance between SD2 and R2N centers of mass. Panel (C) highlights the distance calculated herein between SD2 and R2N centers of mass.

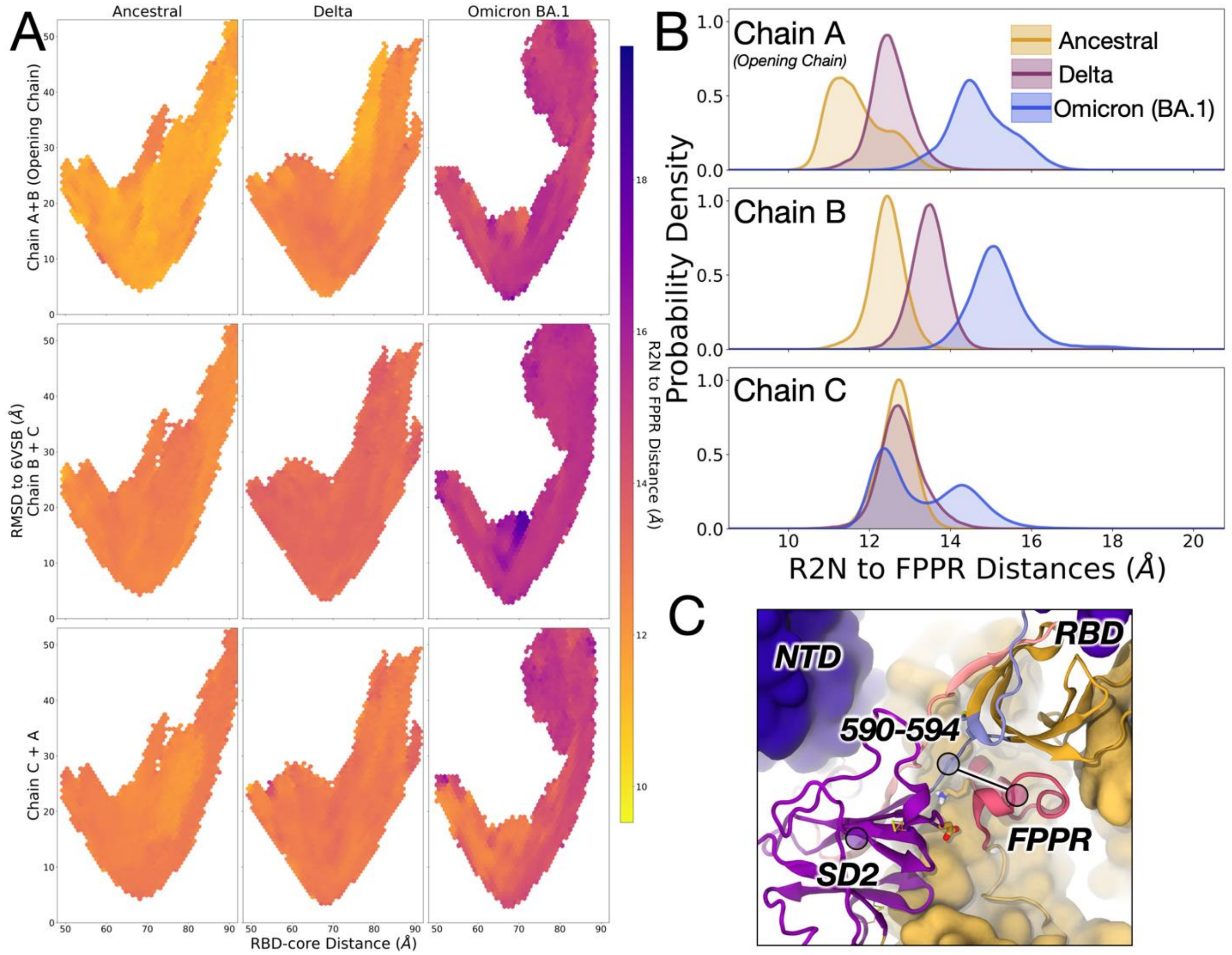


**Figure S11: Distance between the R2N and FPPR centers of mass** for chains A (opening), B, and C, shown as (A) hexagonal histograms as a function of the 2D progress coordinates used to run WE simulations, i.e., RBD-Core Distances (Å) and RMSD to 6VSB (Å), and colored according to the average distance for structures within each bin, and (B) one-dimensional distributions as a function of distance between R2N and FPPR centers of mass. Panel (C) highlights the distance calculated herein between R2N and FPPR centers of mass.

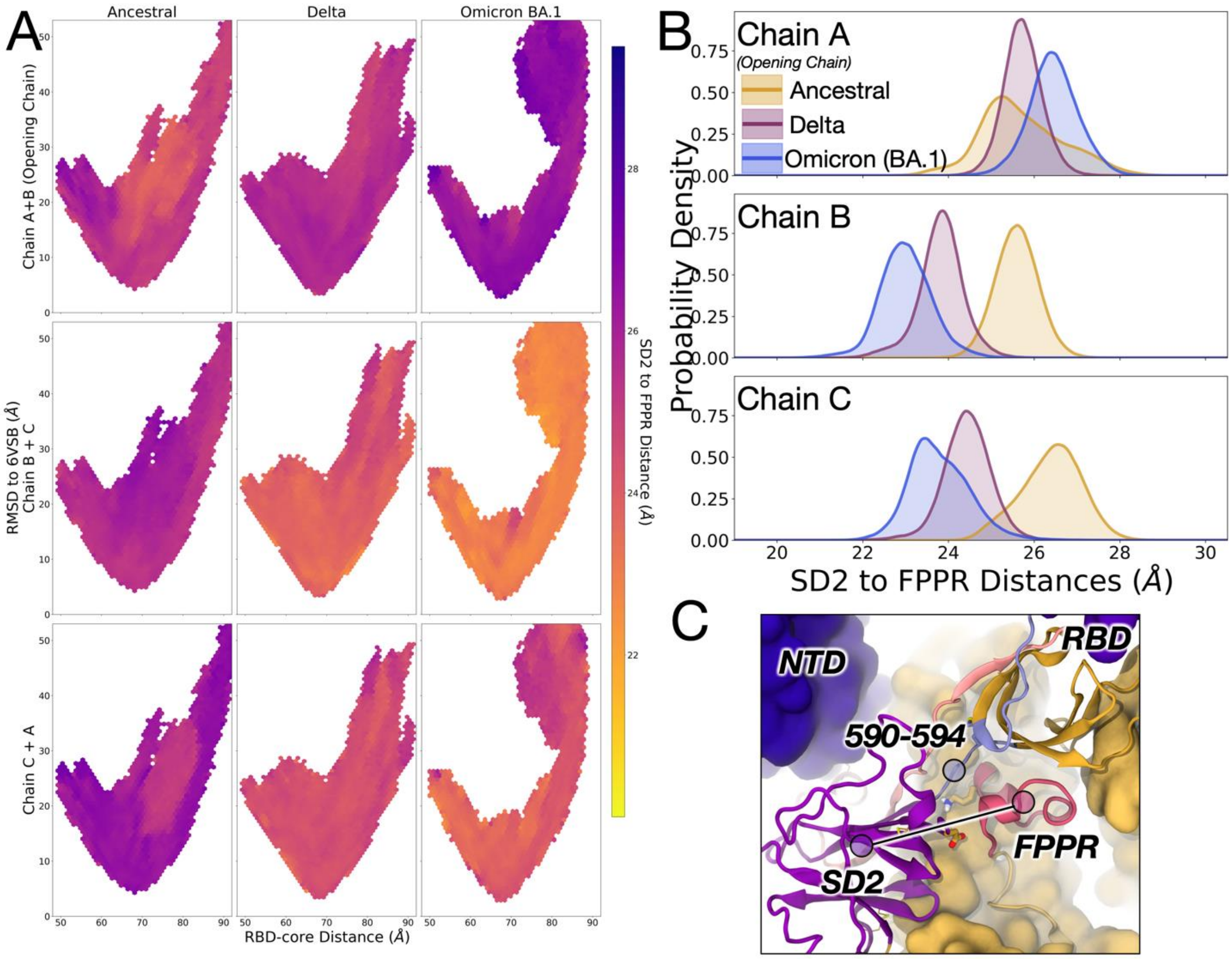


**Figure S12: Distance between the FPPR and SD2 centers of mass** for chains A (opening), B, and C, shown as (A) hexagonal histograms as a function of the 2D progress coordinates used to run WE simulations, i.e., RBD-Core Distances (Å) and RMSD to 6VSB (Å), and colored according to the average distance for structures within each bin, and (B) one-dimensional distributions as a function of distance between FPPR and SD2 centers of mass. Panel (C) highlights the distance calculated herein between FPPR and SD2 centers of mass.

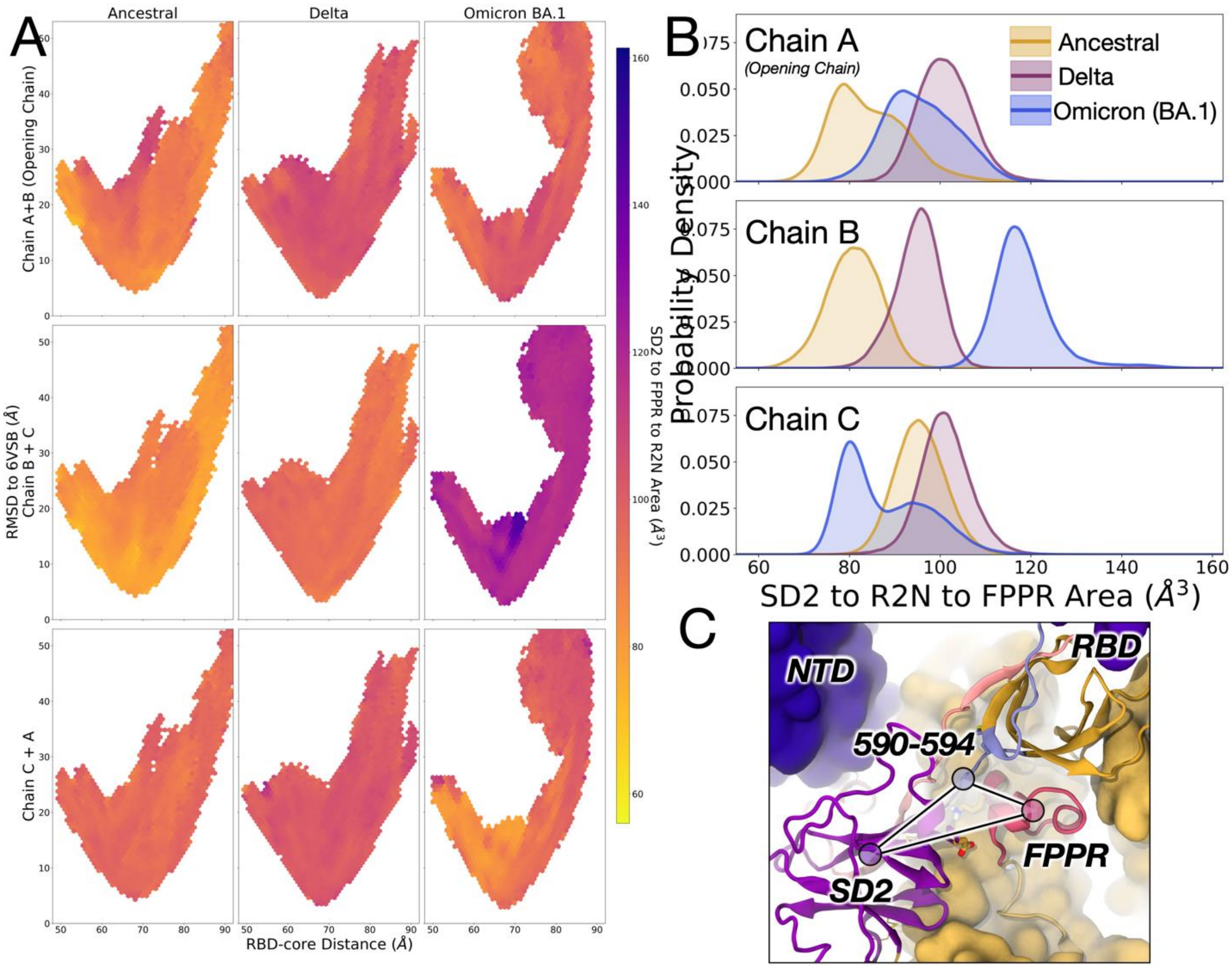


**Figure S13: Area between the SD2, R2N, and FPPR centers of mass** for chains A (opening), B, and C, shown as (A) hexagonal histograms as a function of the 2D progress coordinates used to run WE simulations, i.e., RBD-Core Distances (Å) and RMSD to 6VSB (Å), and colored according to the average distance for structures within each bin, and (B) one-dimensional distributions as a function of distance between FPPR and SD2 centers of mass. Panel (C) highlights the distance calculated herein between FPPR and SD2 centers of mass.

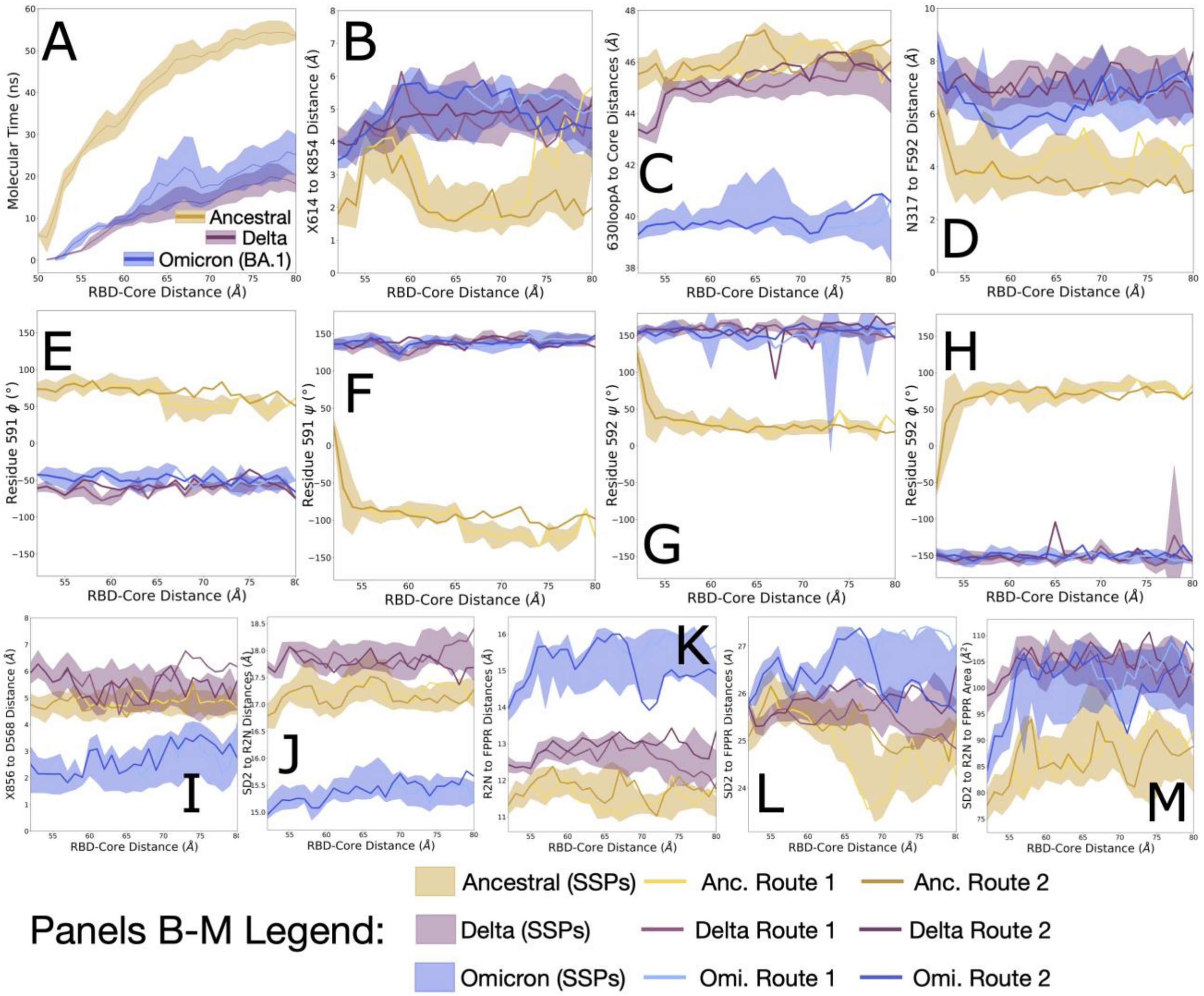


**Figure S14: Trends correlating with spike opening described in the main text shown for all stitched successful pathways (SSPs) for Ancestral, Delta, and Omicron spikes.** Panels B-K also include data collected per clustered Routes 1 and 2. (A) RBD-Core Distance (Å) as a function of average molecular time per stitched successful trajectory. (B) Distance between $X614_A$ to $K854_B$ (Å) as a function of RBD-Core Distance (Å). (C) Distance between $630loop_A$'s center of mass to the spike core's center of mass (Å) as a function of RBD-Core Distance (Å). (D) Distance between $N317_A$ to $F592_A$ (Å) as a function of RBD-Core Distance (Å). (E) Residue $S591_A$'s backbone φ angle (degrees) plotted as a function of RBD-Core Distance (Å). (F) Residue $S591_A$'s backbone ψ angle (degrees) plotted as a function of RBD-Core Distance (Å). (G) Residue $F592_A$'s backbone φ angle (degrees) plotted as a function of RBD-Core Distance (Å). (H) Residue $F592_A$'s backbone ψ angle (degrees) plotted as a function of RBD-Core Distance (Å). (I) Distance between $X856_B$ to $D568_A$ (Å) as a function of RBD-Core Distance (Å). (J) Distance between $SD2_A$'s center of mass to the $R2N_A$'s center of mass (Å) as a function of RBD-Core Distance (Å). (K) Distance between $R2N_A$'s center of mass to the $FPPR_B$'s center of mass (Å) as a function of RBD-Core Distance (Å). (L) Distance between $SD2_A$'s center of mass to the $FPPR_B$'s center of mass (Å) as a function of RBD-Core Distance (Å). (M) Area between $SD2_A$'s center of mass to the $R2N_A$'s center of mass to the $FPPR_B$'s center of mass ($Å^2$) as a function of RBD-Core Distance (Å). Panel A has a dedicated legend. Panels B-K share the same legend.

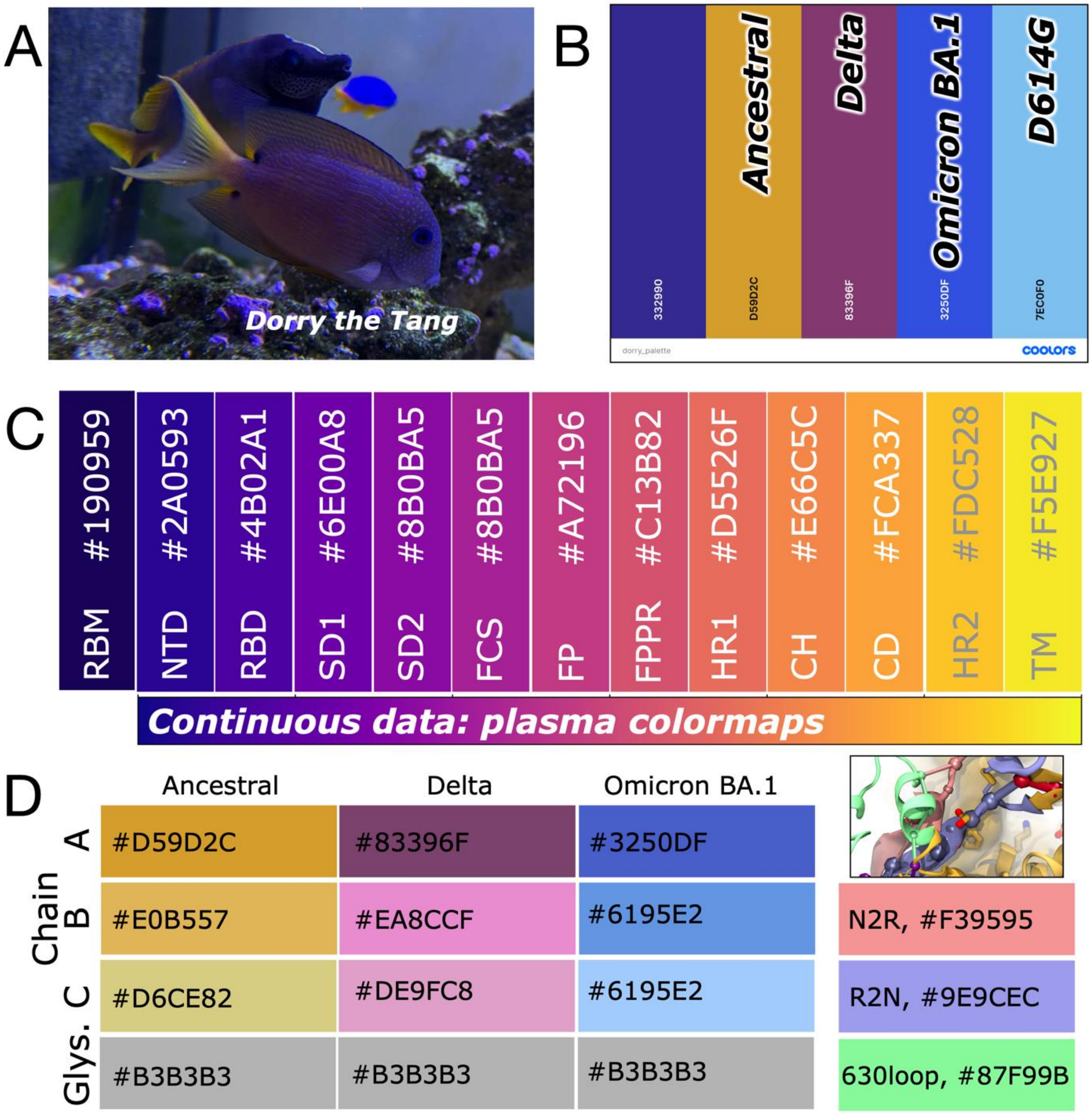


**Figure S15: Color scheme details.** (A) The color scheme used in this work was inspired by the first author's pet fish named Dorry, shown here (photo taken by author). Dorry is a bristletooth tang that lives in the first author's home coral aquarium. (B) Colors chosen from Dorry's image and indication of which colors were used to designate which spike proteins. (C) Continuous data in this work was presented according to one of Matplotlib's colorblind friendly colormaps called plasma. The plasma colormap was also discretized into 14 colors to indicate spike domains. (D) Color codes used to indicate variant spikes and their protomer chains, N- and O-linked glycans, the N2R and R2N linkers, and the 630loop. All images use the font Verdana, a dyslexia friendly font.

(8) Tortorici, M. A.; Beltramello, M.; Lempp, F. A.; Pinto, D.; Dang, H. V.; Rosen, L. E.; McCallum, M.; Bowen, J.; Minola, A.; Jaconi, S.; Zatta, F.; De Marco, A.; Guarino, B.; Bianchi, S.; Lauron, E. J.; Tucker, H.; Zhou, J.; Peter, A.; Havenar-Daughton, C.; Wojcechowskyj, J. A.; Case, J. B.; Chen, R. E.; Kaiser, H.; Montiel-Ruiz, M.; Meury, M.; Czudnochowski, N.; Spreafico, R.; Dillen, J.; Ng, C.; Sprugasci, N.; Culap, K.; Benigni, F.; Abdelnabi, R.; Foo, S.-Y. C.; Schmid, M. A.; Cameroni, E.; Riva, A.; Gabrieli, A.; Galli, M.; Pizzuto, M. S.; Neyts, J.; Diamond, M. S.; Virgin, H. W.; Snell, G.; Corti, D.; Fink, K.; Veesler, D. Ultrapotent Human Antibodies Protect against SARS-CoV-2 Challenge via Multiple Mechanisms.

*Science (1979)* **2020**, *370* (6519), 950–957. https://doi.org/10.1126/science.abe3354.

(22) Case, D. A.; Aktulga, H. M.; Belfon, K.; Cerutti, D. S.; Cisneros, G. A.; Cruzeiro, V. W. D.; Forouzesh, N.; Giese, T. J.; Götz, A. W.; Gohlke, H.; Izadi, S.; Kasavajhala, K.; Kaymak, M. C.; King, E.; Kurtzman, T.; Lee, T.-S.; Li, P.; Liu, J.; Luchko, T.; Luo, R.; Manathunga, M.; Machado, M. R.; Nguyen, H. M.; O'Hearn, K. A.; Onufriev, A. V.; Pan, F.; Pantano, S.; Qi, R.; Rahnamoun, A.; Risheh, A.; Schott-Verdugo, S.; Shajan, A.; Swails, J.; Wang, J.; Wei, H.; Wu, X.; Wu, Y.; Zhang, S.; Zhao, S.; Zhu, Q.; Cheatham, T. E.; Roe, D. R.; Roitberg, A.; Simmerling, C.; York, D. M.; Nagan, M. C.; Merz, K. M. AmberTools. *J Chem Inf Model* **2023**, *63* (20), 6183–6191. https://doi.org/10.1021/acs.jcim.3c01153.

(23) Case, D. A.; Aktulga, H. M.; Belfon, K.; Ben-Shalom, I. Y.; Berryman, J. T.; Brozell, S. R.; Cerutti, D. S.; Cheatham, T. E. I.; Cisneros, G. A.; Cruzeiro, V. W. D.; Darden, T. A.; Forouzesh, N.; Ghazimirsaeed, M.; Giambaşu, G.; Giese, T.; Gilson, M. K.; Gohlke, H.; Goetz, A. W.; Harris, J.; Huang, Z.; Izadi, S.; Izmailov, S. A.; Kasavajhala, K.; Kaymak, M. C.; Kovalenko, A.; Kurtzman, T.; Lee, T. S.; Li, P.; Li, Z.; Lin, C.; Liu, J.; Luchko, T.; Luo, M.; Machado, M.; Manathunga, M.; Merz, K. M.; Miao, Y.; Mikhailovskii, O.; Monard, G.; Nguyen, H.; O'Hearn, K. A.; Onufriev, A.; Pan, F.; Pantano, S.; Rahnamoun, A.; Roe, D. R.; Roitberg, A.; Sagui, C.; Schott-Verdugo, S.; Shajan, A.; Shen, J.; Simmerling, C. L.; Skrynnikov, N. R.; Smith, J.; Swails, J.; Walker, J.; Wang, J.; Wang, J.; Wu, X.; Wu, Y.; Xiong, Y.; Xue, Y.; York, D. M.; Zhao, C.; Zhao, Q.; Kollman, P. A. Amber 2024. University of California, San Francisco 2024.
